# Interplay between bacterial clone and plasmid in the spread of antibiotic resistance genes in the gut: lessons from a temporal study in veal calves

**DOI:** 10.1101/2021.06.19.449090

**Authors:** Méril Massot, Pierre Châtre, Bénédicte Condamine, Véronique Métayer, Olivier Clermont, Jean-Yves Madec, Erick Denamur, Marisa Haenni

## Abstract

Intestinal carriage of extended spectrum β-lactamase (ESBL)-producing *Escherichia coli* is a frequent, increasing and worrying phenomenon, but little is known about the molecular scenario and the evolutionary forces at play. We screened 45 veal calves, known to have high prevalence of carriage, for ESBL-producing *E. coli* on 514 rectal swabs (one randomly selected colony per sample) collected over six months. We characterized the bacterial clones and plasmids carrying *bla*_ESBL_ genes with a combination of genotyping methods, whole genome sequencing and conjugation assays. One hundred and seventy-three ESBL-producing *E. coli* isolates [*bla*_CTX-M-1_ (64.7%), *bla*_CTX-M-14_ (33.5%) or *bla*_CTX-M-15_ (1.8%)] were detected, belonging to 32 bacterial clones, mostly of phylogroup A. Calves were colonized successively by different clones with a trend in decreasing carriage. The persistence of a clone in a farm was significantly associated with the number of calves colonized. Despite a high diversity of *E. coli* clones and *bla*_CTX-M_-carrying plasmids, few *bla*_CTX-M_ gene/plasmid/chromosomal background combinations dominated, due to (i) efficient colonization of bacterial clones and/or (ii) successful plasmid spread in various bacterial clones. The scenario ‘clone vs. plasmid spread’ depended on the farm. Thus, epistatic interactions between resistance genes, plasmids and bacterial clones contribute to optimize fitness in specific environments.

## Introduction

The use of extended-spectrum cephalosporins (ESC) in veterinary and human medicine has led to the emergence of extended spectrum β-lactamases (ESBLs), which confer resistance to these molecules. Since the 80’s (1), ESBL-producing *Enterobacteriaceae* have not only widely disseminated in humans but also in food-producing animals (2–4), among which veal calves in Europe constantly harbored high ESBL intestinal rates (5, 6). Several studies have shown that CTX-M enzymes encoded by plasmid-borne genes are mostly responsible for ESC resistance in veal calves (5, 7), with those from the *bla*_CTX-M_ groups 1 and 9 being the most prevalent (6–9). Of particular concern is also the co-occurrence of *bla*_CTX-M_ and *mcr* genes, the latter conferring resistance to colistin, one of the last-resort antibiotics in human medicine. Indeed, the *mcr-1* gene was found at alarming levels in ESBL-producing *E. coli* isolated from feces of diarrheic veal calves (10, 11).

ESBL dynamics in veal calves has been studied on several occasions at farm level (9, 12). In two previous studies, a marked decrease in ESBL prevalence has been reported, over a ten-week period in three Dutch farms (12), and over the six-month period of the fattening process in ten French farms (9). An extremely high ESBL colonization at admission resulted from calves fed antimicrobial residues-containing waste milk before the first month, and administration of antibiotic collective treatment at the beginning of the fattening, in anticipation of an outbreak (9, 13, 14). Nonetheless, little attention has been paid to genomics insights at ESBL gene, plasmid or bacterial clone levels, which may inform in more detail on each of their respective contributions to ESBL spread during the fattening process.

Here, we followed up a previous study conducted on 45 calves distributed in three fattening farms, from one week after arrival until discharge to slaughterhouses. The central objectives of the study were to clarify *(i)* the link between early intensity and duration of ESBL-producing *E. coli* fecal excretion, *(ii)* the role of plasmids versus bacterial clones in the spread of ESBL genes and *(iii)* the circulation of ESBL-producing *E. coli* clones among calves and their persistence in farms. In addition, we studied the molecular supports of the *mcr-1* gene within ESBL-producing *E. coli* clones. Since the intestinal microbiota is well-known as the epicenter of ESBL selection and ESBL spread in both humans and farm animals (15, 16), the new information reported here in veal calves also contributes to a better understanding of ESBL intestinal spread in general, notably in humans.

## Material and method

### Study design and animal selection

The study was performed in three veal farms (A, B and C) located in the region of Brittany, France. Seven days after their arrival in farms, 50 randomly selected calves per farm were screened for intestinal carriage of ESBL-producing *E. coli.* Rectal swabs were collected and streaked on selective ChromID ESBL agar (bioMérieux, Marcy l’Etoile, France). Calves were classified as “high-level (HL) ESBL carrier”, “low-level (LL) ESBL carrier” or “not (NO) ESBL carrier” based on the number of colonies that grew on the selective medium after 24 hours at 37°C (> 100 colonies= HL, < 100 colonies= LL, or no colony= NO). In each farm, 15 (five of each HL, LL and NO status) out of the initially 50 tested calves were selected and included in the study. Calves were grouped in pens according to their level of ESBL carriage.

### Sampling and antibiotic data collection

Rectal swabs from each of the 15 calves were collected in duplicate every 15 days until departure to the slaughterhouse (Fig. 1). Swabs were placed immediately in portable coolers with ice packs, shipped to the ANSES laboratory in Lyon, France, and stored at −80°C. Antibiotic treatments were recorded by the three farmers throughout fattening (Fig. 1, see Supplementary Methods).

**Fig. 1.**
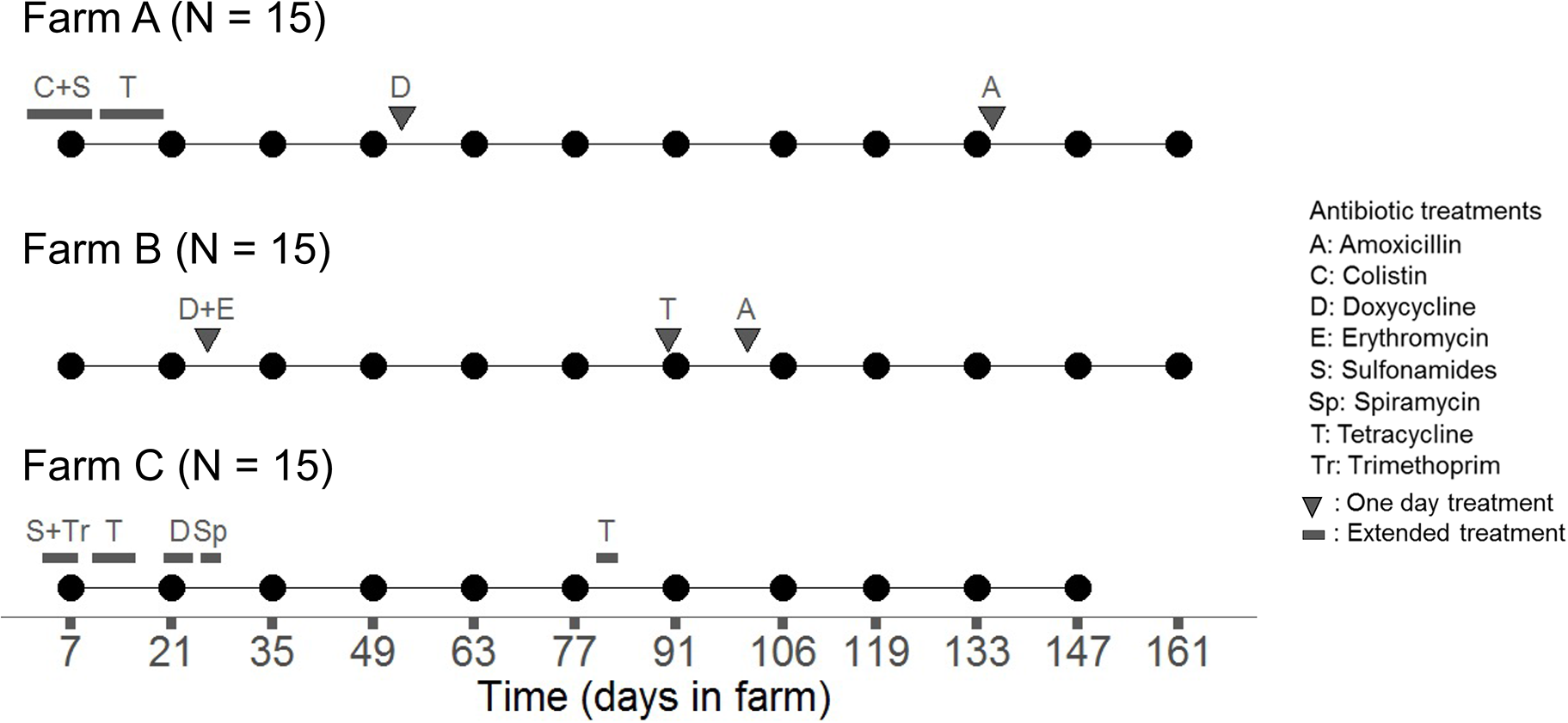
Scheme of sampling dates and collective antibiotic treatments for each veal farm. Sampling points for farm A, farm B, and farm C are represented in the upper panel, middle panel, and lower panel, respectively. “N” indicates the number of calves studied on each farm. The samplings are indicated by black dots. Antibiotic treatments are indicated by bold lines or triangles, and the names of the antibiotics are given in the legend. Of note, one calf on farm C died during the fattening period and was excluded from the study.

### Quantification of *bla*_CTX-M_ copy number in fecal samples

DNA was extracted from swabs collected 7 days and 21 days after arrival in farms, and one by month for each calf, as described in (17), using the DNEasy PowerSoil kit (QIAGEN, Venlo, Netherlands). Quantification of *bla*_CTX-M_ group 1 and group 9 gene copies was performed by quantitative PCR (qPCR) as described in (18), using primers obtained from (19) (see Supplementary Methods). PCR conditions were as described in (19). Products were detected with a LightCycler® 480 System (Roche, Bâle, Switzerland).

### Selection of ESBL-producing *E. coli* isolates

Upon arrival at ANSES, swabs were plated on selective ChromID ESBL agar for the detection of ESBL-producing *E. coli*. Samples were classified as ESBL positive if at least one colony had grown on chromID ESBL agar after incubation at 37°C for 24 hours. One presumptive *E. coli* colony was randomly selected from each selective chromID ESBL plate and stored at −80°C for further characterization.

### Antimicrobial susceptibility testing of ESBL-producing *E. coli* isolates

Antimicrobial susceptibility was tested on all putative ESBL-producing *E. coli* by the disk diffusion method on Mueller-Hinton agar and interpreted according to the clinical breakpoints recommended by the Antibiogram Committee of the French Society of Microbiology (see Supplementary Methods). Susceptibility to ten non-β-lactam antibiotics of veterinary and human interests was tested (colistin, tetracycline, kanamycin, gentamicin, streptomycin, florfenicol, sulfonamides, trimethoprim, nalidixic acid, and enrofloxacin). Minimum inhibitory concentration (MIC) for colistin was determined by the broth microdilution method as recommended by the European Committee on Antimicrobial Susceptibility Testing (EUCAST). For each ESBL-producing isolate, an antibiotic co-resistance score was computed. It was defined as the number of antibiotic classes to which the clone was resistant to (in addition to β-lactamines), among the seven tested (polymyxins, tetracyclines, aminoglycosides, amphenicols, sulfonamides, diaminopyrimidines, quinolones).

### Genotypic discrimination of ESBL-producing *E. coli* isolates

Pulsed-field gel electrophoresis (PFGE) was performed on all putative ESBL-producing *E. coli* isolates using the restriction enzyme *Xba*I. DNA fingerprints were analyzed and the dendrogram of patterns was made using the Dice correlation coefficient, with tolerance and optimization set at 0.5% and 1%, respectively (BioNumerics, Ghent, Belgium). Comparison of PFGE profiles was done among isolates from the same farm to discriminate between ESBL-producing *E. coli* strains circulating among calves (see Supplementary Methods). Phylogenetic grouping was performed using the method described in (20). Detection of *bla*_CTX-M_ group 1, *bla*_CTX-M_ group 9, and *mcr-1* gene was done by PCR as described in (9, 21).

### Whole genome sequencing (WGS) of ESBL-producing *E. coli* isolates

#### Selection of E. coli isolates to be sequenced

We sequenced the genome of a subset of isolates (n=43) based on their PFGE profile. Two isolates were considered different strains if their profiles differed from at least one band. When several isolates had an identical PFGE profile, one isolate was selected for WGS. Isolates having identical PFGE profiles but a different status regarding the detection of the *mcr-1* gene compared to their group were additionally selected for genome sequencing. Genomic DNA was extracted from colonies growing on LBA using the NucleoMag Tissue kit (Macherey-Nagel, Düren, Germany). Libraries were prepared using Nextera DNA library prep kit (Illumina, San Diego, California) and sequenced on an Illumina HiSeq 4000 to produce paired-end reads of 100 base pairs (bp).

#### Phylogenetic analyses and clone definitions

MLST (Achtman and Pasteur Institute schemes), serotype and *fimH* gene allele were determined using SRST2 0.2.0 with standard parameters (22), after an initial quality check (see Supplementary Methods). Genomes were assembled with SPAdes 3.11.1 with the “careful” option to reduce the number of mismatches and short indels. The phylogroup was determined with the *in silico* PCR ClermonTyper 1.4 (23, 24). A core genome was created with ParSNP from Harvest (25–28) and BEDTools (29), using the *E. coli* strain ED1a genome as a reference.

The clone definition was established following the strategy described below. First, isolates were grouped by their haplogroup, which was defined as a combination of their sequence types (ST) according to the Achtman scheme and the Pasteur Institute scheme, their serotype (O:H) and their *fimH* allele. Second, the SNPs detected in the genes of the core genome were used as a genetic distance between the isolates within identical haplogroups. Isolates were considered to be part of the same clone if they had the same phylogroup and haplogroup, and if the number of SNPs in their core genomes was smaller than 100 SNPs. If two isolates had the same phylogroup and haplogroup but their core genome differed by more than 100 SNPs, they were named ‘clone1’ and ‘clone2’ (see Supplementary Methods, Fig. S1). An analysis of the number of SNPs among haplogroups was conducted to (1) confirm the validity of our genotypic markers and to (2) delineate clones (see Supplementary Methods). For the clarity of the reading, clone names are restricted in the text by a combination of their ST and serotype, while their full name (‘phylogroup STs serotype and *fimH* gene allele’) is provided in the figures. The full list and characteristics of the strains sequenced are presented in Table S1A, along with their genome accession numbers.

#### Resistome and virulome analyses

The resistome and virulome of the clones were characterized using the software Abricate 0.8.1 (see Supplementary Methods) (30).

#### Classification of clones according to colonization efficiency

In each farm, clones were grouped according to the number of calves in which they were detected: “inefficient colonizers” when found in 20% of calves or less (corresponding to three calves or less), “intermediate colonizers” when found between 20% and 80% of the calves (four to eleven calves), and “efficient colonizers” when found in 80% of the calves or more (twelve calves or more).

### Conjugation and plasmid sequencing

#### Selection of plasmids to be sequenced

Transfers of plasmids from sequenced isolates into the *E. coli* J53 plasmid-free strain were conducted to characterize the *bla*_CTX-M_ and *mcr-1* carrying plasmids. Conjugation was performed in liquid medium using rifampicin and cefotaxime (5mg/L) or colistin (2mg/L) to select for transconjugants (TC). Only TCs carrying the appropriate plasmid were further characterized (see Supplementary Methods).

#### Sequencing and bioinformatics analyses

Libraries were prepared as described above and sequenced on an Illumina HiSeq 4000 to produce paired-end reads of 100 base pairs (bp). Plasmids were identified by Abricate, using the PlasmidFinder database (30–32). Contigs were sorted according to their chromosomal or plasmidic location using PlaScope (33). The transconjugant reads were mapped on the J53 assembly genome using BWA (34). The unmapped reads were considered as plasmidic and were retrieved with SAMtools 0.1.18 (35, 36). These reads were assembled with SPAdes and the assembled genomes were blasted against the MaGe database (37). Contigs that couldn’t be assembled were blasted against the Refseq database (38). The plasmidic genomes were annotated by RAST (39–41) and the pMLST sequence types were obtained with pMLST-2.0 Server on the CGE. The SNPs were detected by SAMtools (35, 36) and bcltools (42).

#### Phylogenetic analyses and plasmid lineage definition

Plasmid classification was established in a two-step strategy. First, sequenced plasmids were grouped by their incompatibility group, FAB formula for IncF plasmids or ST for the other plasmids, according to the published nomenclature (43–45). Second, the presence of distinct lineages was searched within each group of plasmids. The core genome of a plasmid group was defined as the set of genes present in the genome of the reference and in all the reconstructed genomes. The numbers of SNPs detected in the genes of the core genome were used as a genetic distance to discriminate between plasmid lineages (see Supplementary Methods, Fig. S2). The full list and characteristics of the transconjugants sequenced are presented in Table S1B, along with their genome accession numbers.

#### Identification of bla_CTX-M_ gene carrying-plasmids and mcr-1 gene carrying-plasmids in the collection of isolates

For all isolates, we looked for the plasmid that was identified in the TC of their clone as the one carrying the *bla*_CTX-M_ gene and/or the *mcr-1* gene (see Supplementary Methods for a detailed review of the collection). This was performed by replicon typing (44) using a commercially available kit (Diatheva, Cartoceto, Italy). When needed, the discriminant allele of the pMLST scheme was sequenced (FII allele in the FAB formula, *ardA* gene in the IncI1 pMLST scheme, Genewiz, Leipzig, Germany). Each time the *bla*_CTX-M_ carrying plasmid identified in a clone was detected in its PFGE-related isolates, we hypothesized that the *bla*_CTX-M_ gene was carried by this same plasmid.

### Statistical analyses

We used the Wilcoxon test to compare the proportion of ESBL-positive calves between the first and the last sampling. We used linear regression analysis with 1,000 permutations to test the effect of the level of excretion of ESBL-producing *E. coli* at day 7 on the number of positive samples during the fattening (see Supplementary Methods).

We searched for an association between the total number of copies of *bla*_CTX-M_ genes / g of feces estimated by qPCR and the level of excretion of ESBL-producing *E. coli* (no excretion, low-level, high-level) using Kruskal-Wallis test. This test was also used to search for an association between the number of copies of *bla*_CTX-M_ genes / g and the farm. Post-hoc Dunn’s tests were used for pairwise comparisons between farms, using the Bonferroni method to correct the p-values.

The Spearman correlation test was used to look for an association between the number of calves colonized by a clone and its period of detection in a farm. It was also used to look for an association between these two variables and the antibiotic co-resistance score of each clone. A genome wide association study (GWAS) was done to search for an association between gene content and the ability for clones to be efficient colonizers, or to persist for more than a month in a farm (see Supplementary Methods).

Means are presented with standard deviations for continuous variables and percentages with counts are given for categorical variables. Statistical analyses were performed using R software (R version 3.6.1) (46). The linear regression with permutations was done using the function “lmp” from the package lmPerm (47). Figures were produced using the package ggplot2 (version 2.2.1) (48).

## Results

### Animal inclusion and follow-up

In farm A, the 50 calves screened at day 7 were ESBL positive and 78% (39/50) were HL carriers. Hence, 15 positive calves were included, of which 11 HL carriers. In farm B, 60% [45.2; 73.6]_95%_ of calves (30/50) were ESBL positive at day 7 so that ten ESBL positive and five ESBL negative calves were included, as planned. In farm C, 86% [73.3; 94.2]_95%_ of calves (43/50) were ESBL positive. A switch of calf IDs between LL and NO carrier occurred, which led to the inclusion of 11 ESBL positive calves (five HL and six LL carriers) and four NO carriers. Calves were present for 161 days on farms A and B and 147 days on farm C (Fig.1). One ESBL negative calf from farm C died during fattening and was excluded from the study. There was no missing sample for the 44 remaining calves so that downstream analyses were performed on 514 samples.

### Excretion of ESBL-producing *E. coli* and of *bla*_CTX-M_ genes

There was a significant decrease of ESBL-producing *E. coli* prevalence between the first and the last samplings, five months later (Wilcoxon test, p= 5 x 10^-11^). ESBL-producing *E. coli* were detected in all calves at at least one time point, except for the five ESBL negative calves in farm B, which remained negative over the whole fattening period (Fig. 2A). We thus collected 174 ESBL positive samples (33.9% of the total number of samples), with 84, 15 and 75 positive samples in farms A, B, and C, respectively (Fig. 2A). Calves had an average of 5.6 (± 1.2), 1.0 (± 0.9) and 5.4 (± 1.5) positive samples in farms A, B and C, respectively. Calves that were HL, LL or NO carriers at day 7 had an average of 4.9 (± 2.1), 4.1 (± 2.3) and 2.0 (± 1.4) positive samples, respectively. No difference in the number of positive samples was evidenced between calves with HL or LL excretion at day 7 (linear regression with 1,000 permutations, p= 0.08, see Supplementary Results). NO carrier calves at day 7 had significantly fewer positive samples than HL and LL carriers at day 7 (linear regression with 1,000 permutations, p= 7 x 10^-5^ and p= 0.003, respectively).

**Fig. 2.**
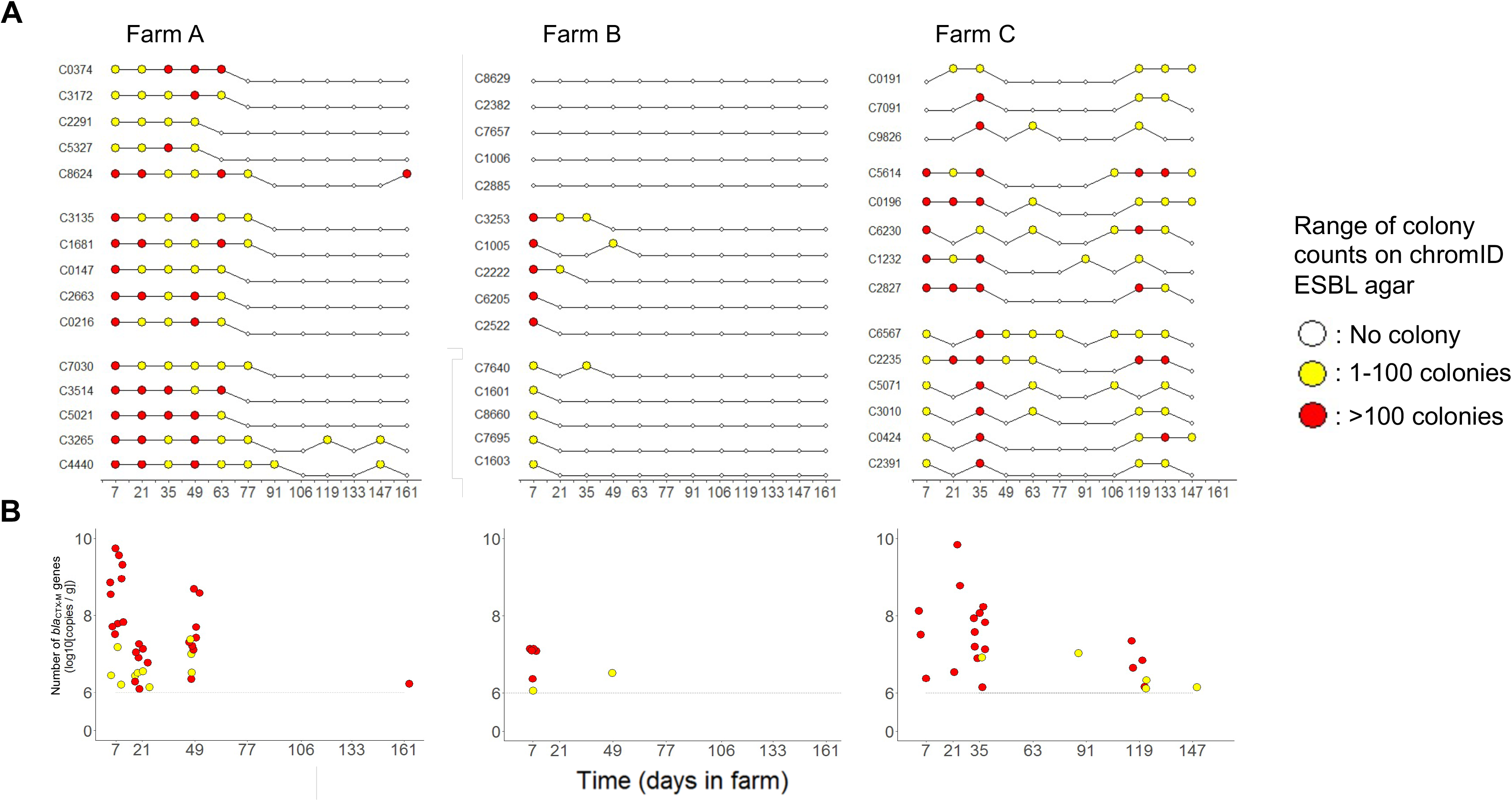
Quantification of ESBL-producing *Enterobacteriaceae* carriage and of *bla*_CTX-M_ gene copy numbers. For each sample, panel A represents the load of colonies on ESBL-producing *Enterobacteriaceae* selective medium after 24 hours at 37°C. Samples are represented by dots and are arranged according to farms, calves, and time. Dots linked by a line are the samples of the same calf, which ID is provided on the left of each line. Calves of farm A, farm B, and farm C are represented on the left, middle, and right of the panel A, respectively. Calves are clustered according to the pen they shared. Panel B represents the total number of copies of *bla*_CTX-M_ genes estimated by qPCR targeting group 1 and group 9. The limit of quantification was 10^6^ copies of *bla*_CTX-M_ genes / g of feces (indicated by a dotted line). Each dot represents one sample. For each calf, quantification was done on samples collected 7 days and 21 days after arrival in the farm, and on one sample by month. The days corresponding to samples processed are indicated on the x-axis. Samples of farm A, farm B, and farm C are represented on the left, middle, and right of panel B, respectively.

The *bla*_CTX-M_ genes were detected in 68 samples among the 307 samples tested by qPCR (Fig. 2B). The number of colonies on ChromID ESBL agar was associated with the number of copies of *bla*_CTX-M_ / g of feces estimated by qPCR (Kruskal-Wallis test, p< 10^-15^, Table S2). For 73.8% of the samples classified as LL (48/65), the number of copies of *bla*_CTX-M_ / g was below 10^6^, while for 89.3% of the samples classified as HL (50/56), the number of *bla*_CTX-M_ / g was above 10^6^ (Fig. 2B). The number of copies of *bla*_CTX-M_ / g of feces was significantly different between farms (Kruskal-Wallis test, p= 8 x 10^-8^), and was significantly higher in farms A and C compared to farm B (Dunn tests, Bonferroni corrected p= 2 x 10^-7^ and p= 7 x 10^-5^, respectively).

### Characterization of ESBL-producing *E. coli* isolates

One colony was isolated from the 174 positive samples for further characterization. In farm C, one isolate was lost (isolated at day 21 from the calf ‘C0191’). Thus, downstream analyses were conducted on 173 isolates. The ESBL-producing *E. coli* isolates carried either *bla*_CTX-M-1_ (112/173 isolates, 64.7%), *bla*_CTX-M-14_ (58/173, 33.5%) or *bla*_CTX-M-15_ (3/173, 1.8%). All of them were resistant to at least two antimicrobials other than ß-lactams, the most common resistances being against tetracyclines, sulfonamides, trimethoprim and aminoglycosides (Table S3A).

The 173 isolates were grouped in 43 different combinations (Fig. S3A, Fig. S3B, Fig. S3C). The genome of one representative isolate of each of the 43 combinations was sequenced (see Supplementary Results). As defined in the material & method section, 32 *E. coli* clones were discriminated among the 43 isolates sequenced. A high phylogenetic diversity was observed among ESBL-producing *E. coli* clones, as clones from all phylogroups except B2 were detected, and one clone had an unassigned phylogroup (‘ST1850/918 O9:H10’, Fig. S4A). Of note, the ST58 lineage from the B1 phylogroup, known as part of the clonal complex 87 (CC87) (49), was detected in farms B and C, with the isolation of the two clones ‘ST58/24 O8:25’ and ‘ST58/24 O9:H25’ (Fig. 3). ESBL-producing *E. coli* clones had a mean co-resistance score of 4.8 (± 1.0), the minimum and maximum being two and seven, respectively.

**Fig. 3.**
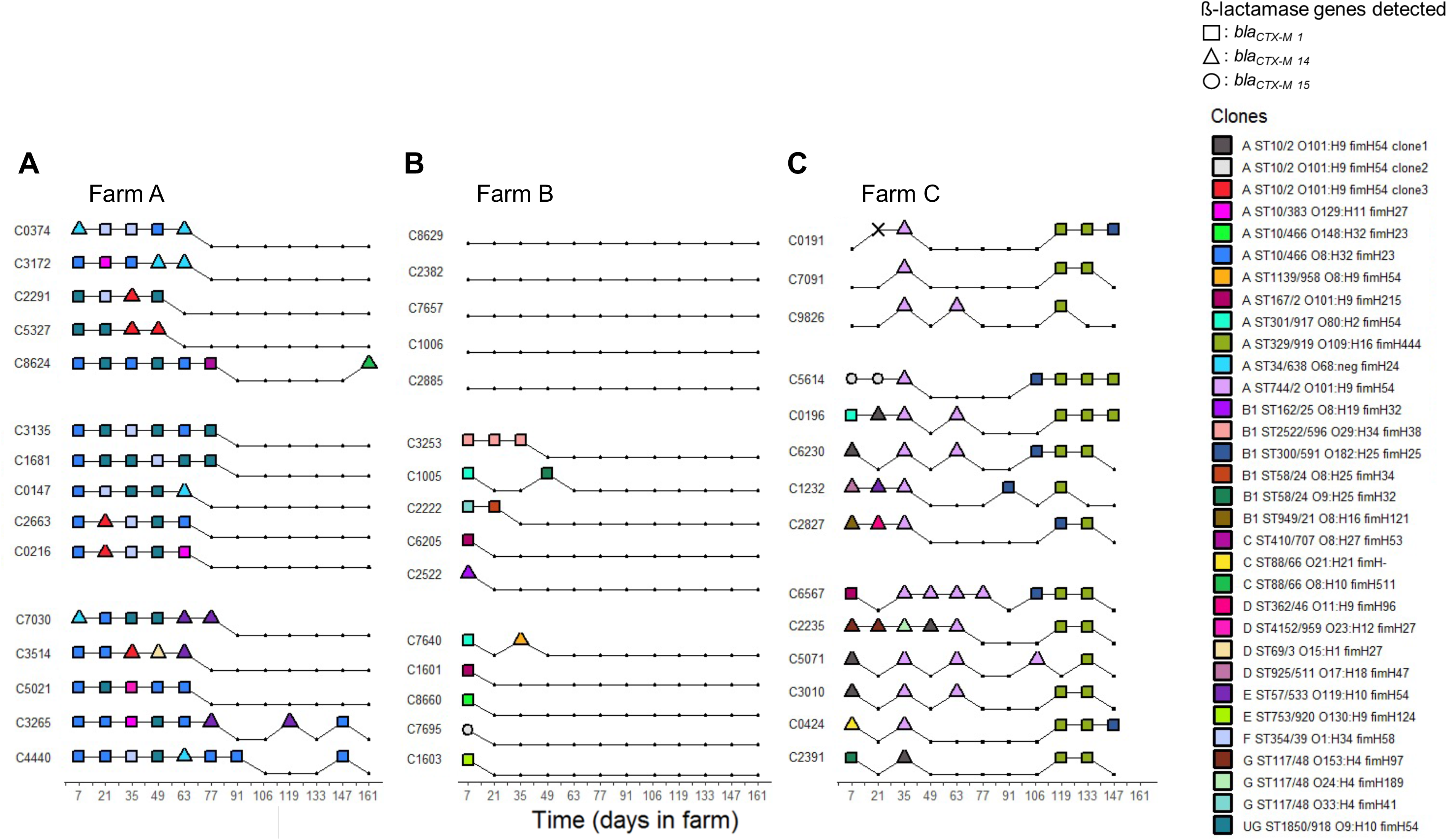
Detection of ESBL-producing *E. coli* clones in each farm over the fattening. For each sample, one ESBL-producing *E. coli* isolate was randomly selected for molecular characterization. ESBL-producing *E. coli* isolates were grouped according to their PFGE profiles. At least one isolate per group of identical PFGE profiles was selected for WGS. A clone was defined according to the combination of the phylogroup, ST Achtman/ST Pasteur Institute, serotype, and *fimH* allele gene identified. Two isolates harboring the same PFGE profile were considered to be related to the same clone. The distribution of the clones detected in farms A, B, and C are represented on panels A, B, and C, respectively. The days are indicated on the x-axis of each panel. Calves are clustered according to the pen they shared. Dots linked by a line are the samples of the same calf, which ID is provided on the left of each line. The color of the dot indicates the ESBL-producing *E. coli* clone that was isolated from this sample, and its shape represents the *bla*_CTX-M_ gene it carried. Clones’ full names are indicated in the legend. In farm C, one isolate was lost before any genotypic characterization and is depicted by an ‘X’ (isolated at day 21 in calf ‘C0191’). ‘UG’: unassigned phylogroup.

ESBL-producing *E. coli* clones carried in average 9.6 (± 2.5) antibiotic resistance genes (see Supplementary Results), and genes found in more than 50% of the clones conferred resistance against penicillins, tetracyclines, aminoglycosides and sulfonamides (Fig. S4C). These genes were *bla*_TEM-1_ (27/32 clones, 84.4%), *tet(A)* (27/32, 84.4%), *aph*(*6*)*-Id* (25/32, 78.1%)*, aph(3’’)-Ib* (23/32, 71.9%), *sul2* (21/32, 65.6%), *aph(3’)-Ia* (20/32, 62.5%), and *sul1* (17/32, 53.1%). Virulence genes associated with intestinal and extra-intestinal pathogenicity were found in all ESBL-producing clones (Fig. S5A, Fig. S5B and Fig. S5C). ESBL-producing clones carried on average 34 (± 9) genes associated with extraintestinal virulence and 19 (± 14) genes associated with intestinal virulence (Fig. S5A, Fig. S5B).

Three potential intestinal pathogenic clones were found: the clone ‘ST301/917 O80:H2’ was an enterohemorrhagic (EHEC) / extra-intestinal *E. coli* hybrid pathotype, first described in (50), harboring the locus of enterocyte effacement (LEE) with the intimin-encoded *eae*-ξ gene, the *ehxA, efa1, espP* genes, the s*tx2a* and *stx2b* genes (Fig. S5B). It was also carrying *iss, iroN hlyF, ompT,* four genes associated with the pS88 plasmid, which is related to avian pathogenic *E. coli* plasmids (51) (Fig. S5A and Fig. S5B). The clone ‘ST329/919 O109:H16’ was a STEC clone (*stx2a* and *stx2b* genes detected, Fig. S5B). The clone ‘ST300/591 O182:H25’ was an atypical EPEC clone (LEE detected, Fig. S5B).

### Within-farm dynamics of ESBL clones and plasmids

The *bla*_CTX-M-1_ gene was carried by five plasmid types disseminated in 16 clones (encompassing 112/173 isolates, 64.7%, Fig. 4), while the *bla*_CTX-M-14_ gene was carried by five plasmid types disseminated by 15 clones (encompassing 58/173 isolates, 33.5%, Fig. 5). The *bla*_CTX-M-1_ and *bla*_CTX-M-14_ genes were carried by distinct plasmids and clones (see Supplementary Methods for details). Of note, the *bla*_CTX-M-15_ gene was detected on the chromosome of the same clone in farms B and C (Fig. 3, Fig. S6A, see Supplementary Results).

**Fig. 4.**
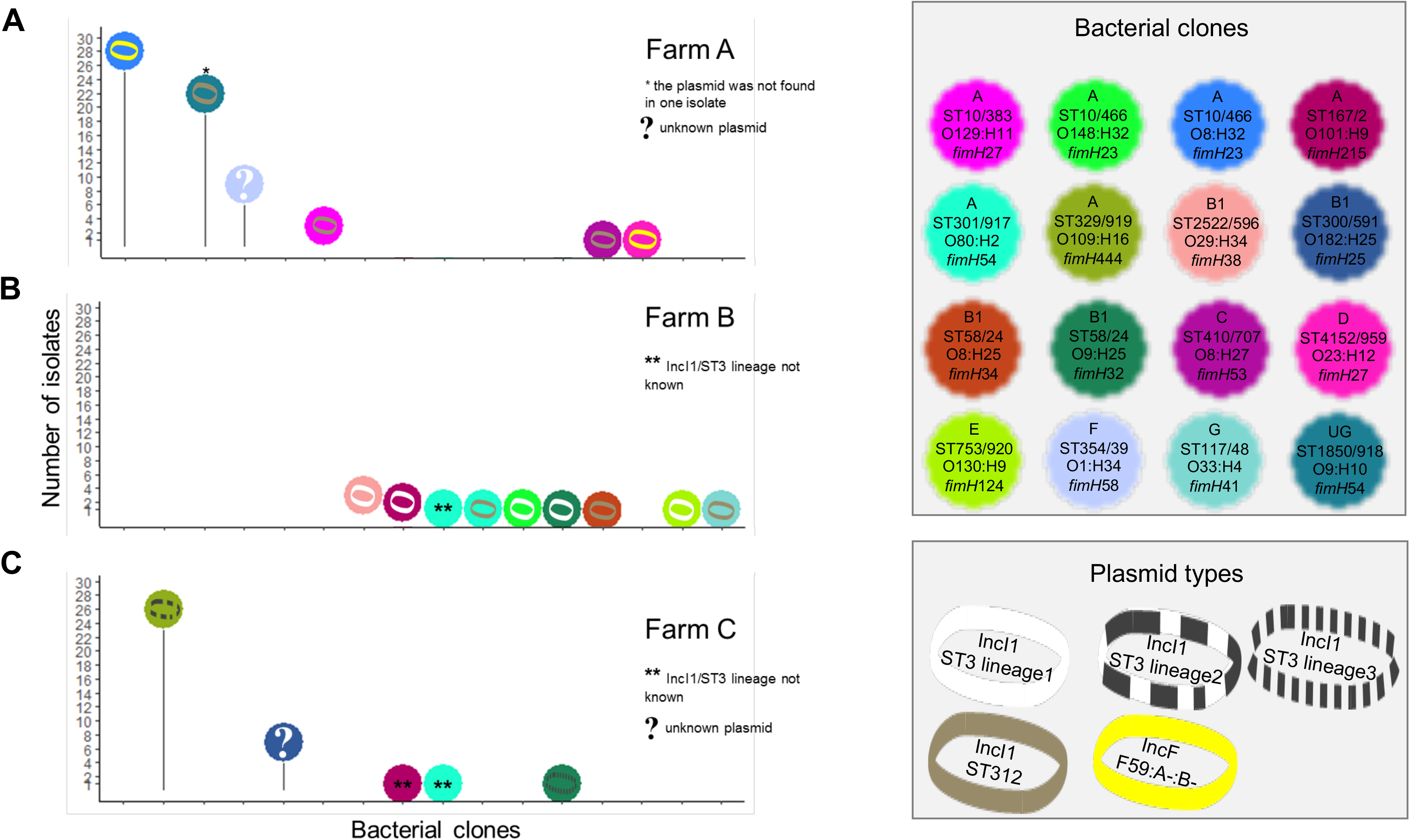
Characterization of *E. coli* clone and plasmid combinations carrying *bla*_CTX-M-1_ gene. Number of isolates for each ESBL-producing *E. coli* clones carrying the *bla*_CTX-M-1_ gene and its molecular supports. The distribution of the clones detected in farms A, B, and C are represented on panels A, B, and C, respectively. Genotyping methods involving PFGE and followed by WGS were used to discriminate bacterial clones. Plasmids carrying a *bla*_CTX-M_ gene were characterized using WGS after conjugation in a K-12 strain. The number of Single Nucleotide Polymorphisms (SNPs) on the plasmid core genome was used to discriminate plasmid lineages, using a threshold of 100 SNPs. For each ESBL-producing *E. coli* clone, we looked for the presence of the plasmid carrying *bla*_CTX-M_ gene in all of its isolates by replicon typing. When needed, the discriminant allele of the pMLST scheme was sequenced (FII allele in the FAB formula, *ardA* gene in the IncI1 pMLST scheme).

**Fig. 5.**
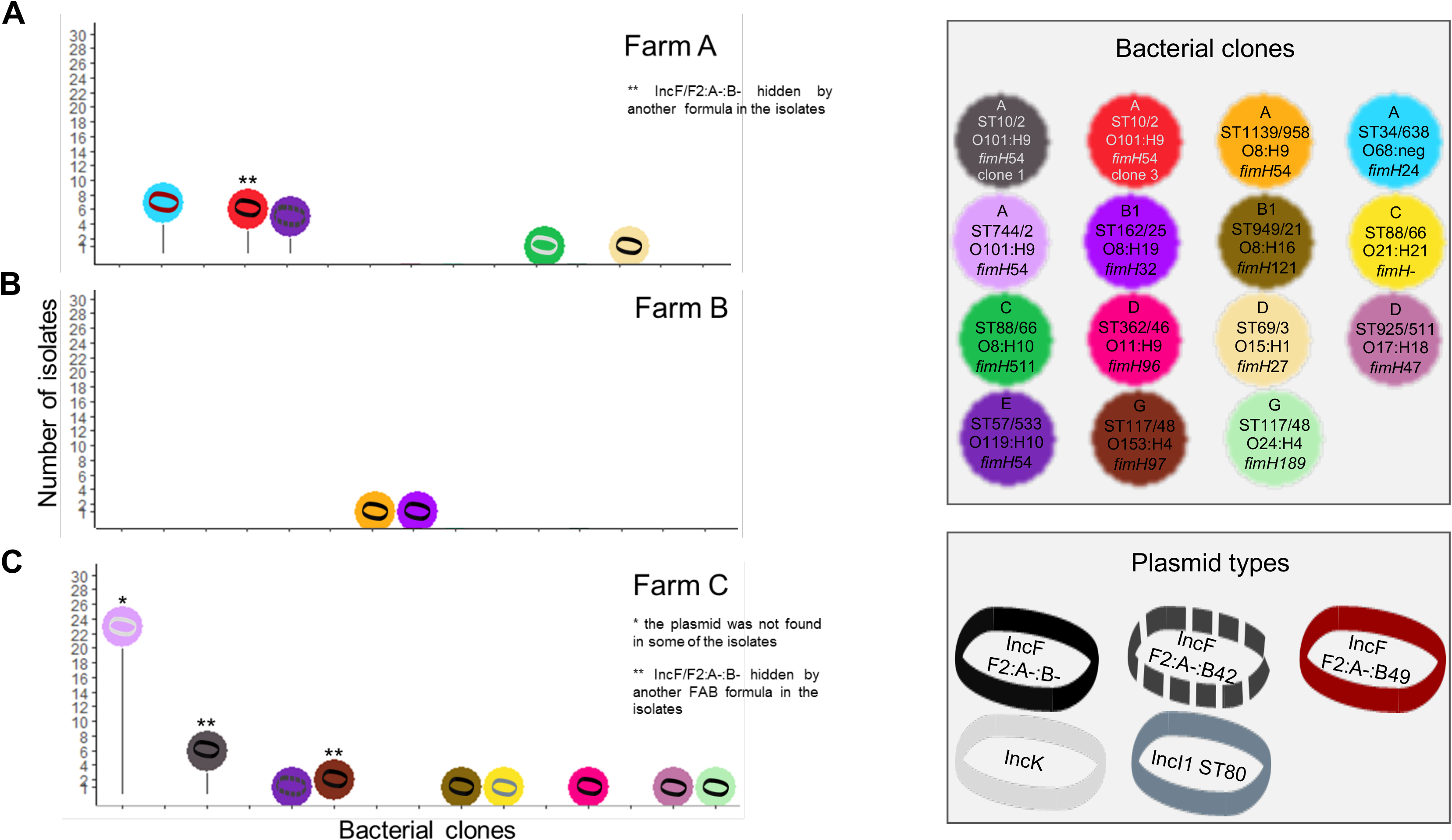
Characterization of *E. coli* clone and plasmid combinations carrying *bla*_CTX-M-14_ gene. Number of isolates for each ESBL-producing *E. coli* clones carrying the *bla*_CTX-M-14_ gene and its molecular supports. The distribution of the clones detected in farms A, B, and C are represented on panels A, B, and C, respectively. Genotyping methods involving PFGE and followed by WGS were used to discriminate bacterial clones. Plasmids carrying a *bla*_CTX-M_ gene were characterized using WGS after conjugation in a K-12 strain. The number of Single Nucleotide Polymorphisms (SNPs) on the plasmid core genome was used to discriminate plasmid lineages, using a threshold of 100 SNPs. For each ESBL-producing *E. coli* clone, we looked for the presence of the plasmid carrying *bla*_CTX-M_ gene in all of its isolates by replicon typing. When needed, the discriminant allele of the pMLST scheme was sequenced (FII allele in the FAB formula, *ardA* gene in the IncI1 pMLST scheme).

In farm A, 11 *E. coli* clones were identified. The mean number of clones colonizing a calf was 3.5 (± 0.7). All calves experienced long-term excretion through a succession of colonizations during the first half of the fattening period (Fig. 3A). Most of these clones had diffused among the batch of calves, with 63.6% of them (7/11) detected in several calves. The *bla*_CTX-M-1_ gene was predominant and carried by at least two plasmids in five clones (encompassing 64/84 isolates in this farm, Fig. 4A). The presence of *bla*_CTX-M-1_ gene was mostly due to two efficient colonizers representing 78.1% (50/64) of the isolates carrying this gene and 59.5% (50/84) of all isolates in this farm. They were widespread among the batch of calves, as more than 80.0% (12/15) of the calves excreted them (Fig. 3A).

In farm B, 11 *E. coli* clones were identified and the mean number of clones colonizing a calf was 1.3 (± 0.5). Calves experienced a short-term excretion, which ended after 49 days (Fig 3B). All *E. coli* clones were inefficient colonizers. The *bla*_CTX-M-1_ gene was predominant (12/15 isolates) and spread by IncI1/ST3 lineage 1 and IncI1/ST312 plasmids distributed in five and three clones, respectively (Fig. 4B). The *bla*_CTX-M-14_ gene was detected once in two calves that had distinct clones, but which had the same *bla*_CTX-M-14_-carrying IncF/F2:A-:B-plasmid (Fig. 5B). In this farm, a higher level of plasmid diffusion than *E. coli* clone diffusion among calves explained the presence of *bla*_CTX-M_ genes.

In farm C, 14 *E. coli* clones were identified and the mean number of clones colonizing a calf was 3.6 (± 1.0). Calves experienced an unsteady excretion, characterized by a succession of ESBL clone colonizations until the end of the fattening period (Fig. 3C). Only 28.6% (4/14) of clones were found in several calves. From the second month until the end, calves were successively colonized by only three *E. coli* clones: ‘ST744/2 O101:H9’ detected from day 35 to day 106, ‘ST300/591 O182:H25’ from day 91 to day 147, and ‘ST329/919 O109:H16’ from day 119 to day 147 (Fig. 3C). The *bla*_CTX-M-1_ and *bla*_CTX-M-14_ genes were both detected in 36 isolates. The presence of *bla*_CTX-M-1_ and *bla*_CTX-M-14_ genes was mostly due to two *E. coli* clones found in 100% (14/14) and 93.8% (13/14) of the calves, respectively (Fig. 3C, see Supplementary Results). These two efficient colonizers each carried different plasmids that were themselves not efficient spreaders, as no other *E. coli* clone in this farm carried these plasmids (Fig. 4C, Fig. 5C).

### Between-farm dynamics of ESBL clones and plasmids

Most *E. coli* clones were found only in one farm, except for five clones (‘ST10/2 O101:H9 clone2’, ‘ST167/2 O101:H9’, ‘ST301/917 O80:H2’ ‘ST58/24 O9:H25’, and ‘ST57/533 O119:H10’, Fig. 6, Fig. S4A). These five clones had been isolated one or two times in no more than three calves in each farm (Fig. 6). Hence, although detected in several farms, these clones were inefficient colonizers in all of them. Four ESBL plasmids were identified in several farms: IncI1/ST312 in farms A and B, IncK in farms A and C, IncF/F2:A-:B-in all three farms, and IncF/F2:A-:B42 in farms A and C (Fig. 4, Fig. 5). IncF/F2:A-:B42 plasmid was spread by the same *E. coli* clone in different farms (‘ST57/533 O119:H10’, Fig. 5). The other plasmids were carried by distinct *E. coli* clones in the different farms. While present in two farms, IncK and IncF/F2:A-:B42 plasmids were detected only in one clone per farm, suggesting a limited spread (Fig. 5). On the opposite, IncI1/ST312 and IncF/F2:A-:B-have spread in several clones in each farm. The IncI1/ST312 plasmid has spread in a few clones, and notably met the efficient colonizer ‘ST1850/918 O9:H10’ in farm A (Fig. 4A). IncF/F2:A-:B-was present in the three farms and has spread in at least two *E. coli* clones in each of them (Fig. 5). In farm C, it has spread in six clones, suggesting higher efficiency to disseminate compared to the other plasmids.

**Fig. 6.**
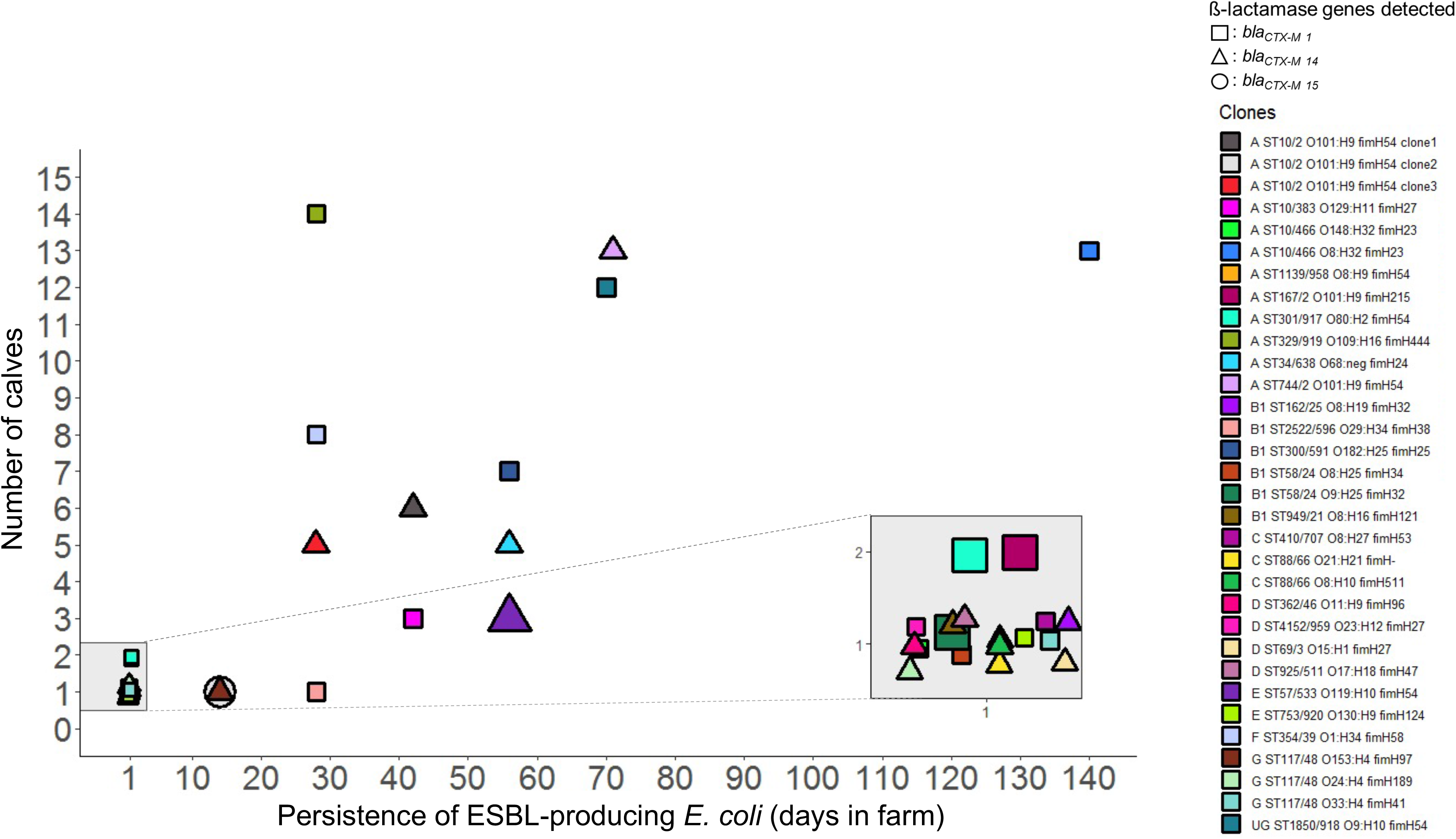
Number of calves colonized by ESBL-producing *E. coli* clones according to their duration of detection in farms. The duration of detection of each clone is represented on the x-axis, and corresponds to the difference between the last day and the first day a clone was detected in a farm. The y-axis represents the number of calves within a farm in which clones were detected. Each dot represents an ESBL-producing *E. coli* clone, and its shape represents the *bla*_CTX-M_ gene it carried. The size of clones found in several farms is increased compared to the clones that were detected in one farm. For clones that were found in several farms, the longest detection time and the highest number of calves colonized is provided. A zoom on the clones found at only one sampling time was added to increase the readability of the figure.

### Persistence of ESBL clones over the fattening period

The length of the period of detection was significantly associated with the number of calves in which ESBL-positive bacterial clones were detected (Spearman’s correlation, p= 4 x 10^-9^). The correlation coefficient was equal to 0.85, indicating a strong positive association between the diffusion of ESBL clones among a batch of calves and their persistence in the farm during fattening. No significant association was found between the antibiotic co-resistance score of the ESBL clones and their diffusion among a batch of calves, nor with their persistence in the farms. No significant association was found between the genome content of the ESBL clones and their diffusion among a batch of calves, nor with their persistence in the farms (GWAS analyses, Bonferroni-corrected p>0.05).

### Distribution of the *mcr-1* gene in bacterial clones and plasmids

The *mcr-1* gene was present in nine clones, eight of which in farm A and one in farm C (encompassing 53 isolates in total, Fig. 7). Calves from farm A received early colistin treatment and were the only ones exposed to colistin during the fattening period (Fig. 1). Plasmids carrying the *mcr-1* gene found in transconjugants were IncX4 and IncHI2 plasmids (Fig. 7). The *mcr-1*-carrying-IncX4 plasmid disseminated in three clones and was present in 73% of the *mcr-1*-positive isolates in farm A. Of note, a chromosomal insertion of the *mcr-1* gene was found in two isolates, one from farm A and one from farm C (Fig. S6A and S6A, see Supplementary Results).

**Fig. 7.**
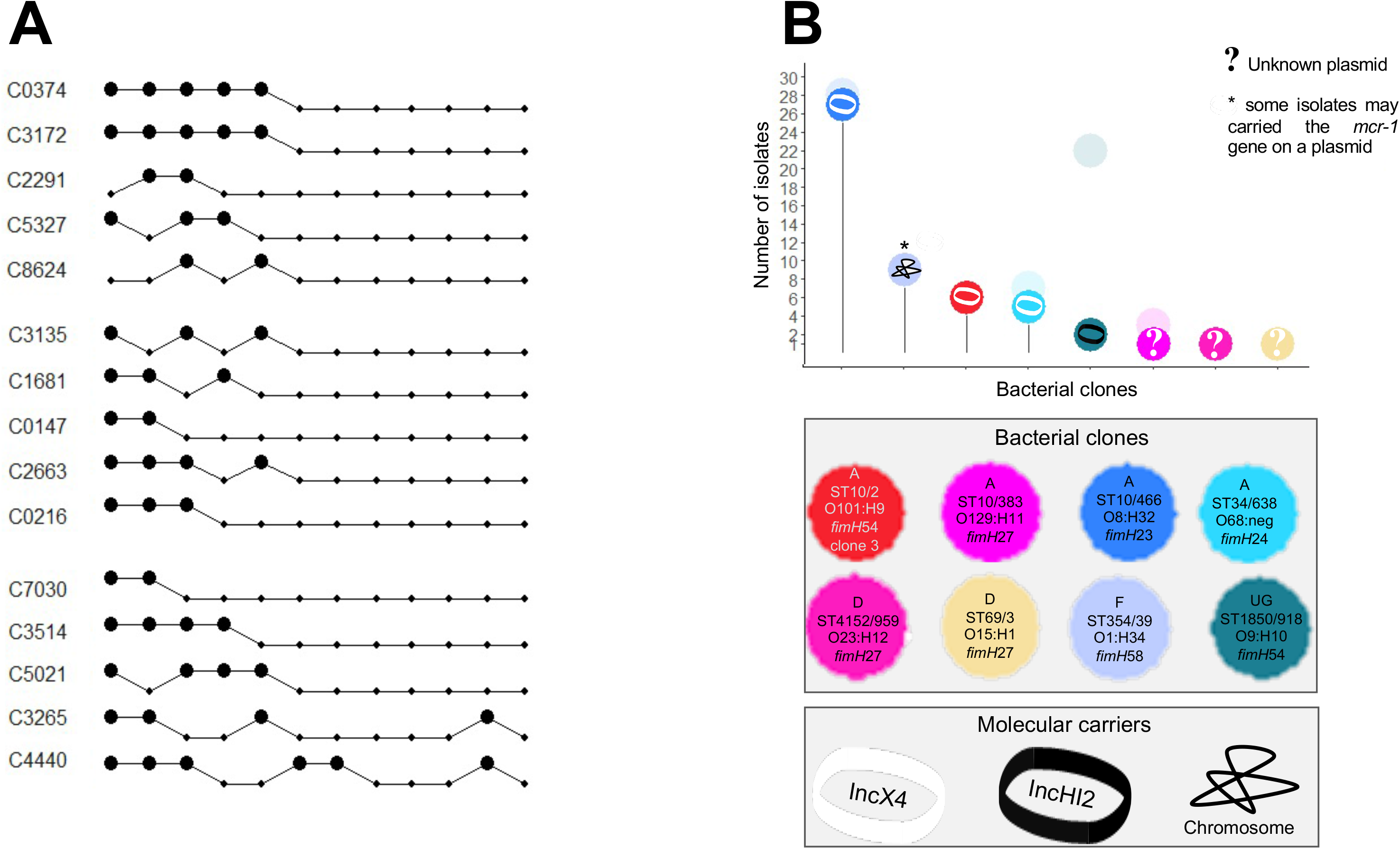
Detection of ESBL-producing *E. coli* clones carrying the *mcr-1* gene and characterization of *mcr-1* gene-carrying *E. coli* clone and plasmid combinations in farm A. Panel A represents samples in which ESBL-producing *E. coli* isolates carrying the *mcr-1* gene (big black dots) in farm A. In each isolate, detection of the *mcr-1* gene was done by PCR. Dots linked by a line are the samples of the same calf, which ID is provided on the left of each line. Calves are clustered according to the pen they shared. Genotyping methods involving PFGE and followed by WGS were used to discriminate bacterial clones. Plasmids carrying the *mcr-1* gene were characterized using WGS after conjugation in a K-12 strain. The number of Single Nucleotide Polymorphisms (SNPs) on the plasmid core genome was used to discriminate plasmid lineages, using a threshold of 100 SNPs. Panel B represents the number of isolates per combination carrying the *mcr-1* gene in farm A. As the *mcr-1* gene was not found in all the isolates of a given ESBL-producing *E. coli* clone, transparent dots representing the total number of isolates of a given clone were added. For each ESBL-producing *E. coli* clone, we looked for the presence of the plasmid carrying the *mcr-1* gene in all of its isolates by replicon typing.

## DISCUSSION

### Dynamics of ESBL prevalence in calves

In accordance with previous studies (9, 12), a decrease in ESBL prevalence in calves was observed over the fattening period, although each farm displayed a specific dynamic scheme (Fig. 2). Differences in ESBL dynamics among farms may be attributed to local factors, such as intercurrent diseases justifying antibiotic treatment. All ESBL positive clones were resistant to several antibiotics commonly used in the veal industry (9, 52–54) (Table S3A, Fig. S4C). Additionally, in farm A, the early exposure to colistin likely selected ESBL positive clones co-harboring the *mcr-1* gene. Of note, whereas ESBL positive clones are usually present in the subdominant flora (6, 9), the opposite situation may have happened in farm A during the first 10 days, reflecting a massive abundance of ESBL positive clones concomitant with ongoing colistin treatment (Fig.1, Fig. 2). Furthermore, in accordance with a previous study, (9) while the prevalence of ESBL-producing *E. coli* decreased over time (Fig. 2), it has been shown that non ESBL-producing *E. coli* remained at high levels (6, 9). These data suggest that, in the absence of ESC treatment in the three farms (Fig.1), maintaining a *bla*_CTX-M_-carrying plasmid could have entailed a cost to the ESBL-producing clones. Such cost, caused by plasmid replication and the expression of plasmid-borne genes (55–57), could alter bacterial host metabolic gene expression, ultimately impairing their ability to maintain a competitive growth rate in an environment with limited resources (58, 59).

Loss of *bla*_CTX-M_ carrying plasmid may have happened during our study, but as we focused only on *E. coli* clones displaying a phenotype of ESBL production, we cannot estimate the magnitude of such loss rate. Studies showed that spontaneous loss happens rarely *in vivo*, even when ESBL-producing strains compete with plasmid-free strains, as shown in pigs, rabbits and lambs (60–63). A modulation of the cost by a selective switch off of horizontally acquired gene expression (64, 65) could also explain why no association was found between genes and the ability for a clone to colonize many calves nor to persist in farms.

### Long-term carriage is associated with recurrent colonizations by ESBL-producing E. coli clones

We showed that *bla*_CTX-M_ gene-long-term carriage was characterized by successive detections of distinct ESBL clones in calves (Fig. 3), which highlights that ESBL carriage in veal farms is a highly dynamic process. These data extend similar observations from our group and others on ESBL clones colonizing the human microbiota, for instance where multiple transient colonizations by up to seven *E. coli* clones was also found to drive the intestinal carriage of *bla*_CTX-M_ genes over a 22-days period (66) in humans visiting high-risk areas for acquiring ESBL-producers (67). Such similar data on intestinal dynamics in ESBL colonization is noticeable considering the differences in gut microbiota composition between humans and cattle at inter-species level (68–70) and intra-species level (71, 72). Moreover, phylogroups B1, E and A, very common in cattle (71–73), accounted for a large proportion of the ESBL clones that proved very diverse, suggesting multiple acquisition events of *bla*_CTX-M_ genes. Of note, phylogroup B2, predominant in human and companion animal’s commensal populations (72, 74) but minor in cow’s populations (71), was the only phylogroup not detected. In humans, ESBL clones of phylogroups B2, D and F rather than A, B1 and E persisted longer in travelers returning from tropical regions (Armand-Lefevre *et al*., Microbial Genomics, in press). Similarly, here in calves, alongside acquisition events, the successful spread of ESBL genes in gut microbiomes is fostered by chronic recurrent intestinal colonization by non-B2 bacterial hosts, *i.e.* that are well-adapted to the bovine host. Moreover, we found that the number of copies of *bla*_CTX-M_ / g of feces was significantly higher in farms A and C compared to farm B (Fig. 2), suggesting that ESBL producers’ abundance at start could help predict the persistence of ESBL producers’ carriage, as it was shown in travelers (67).

### Efficient bla_CTX-M_ spread is supported by a few E. coli clones

In two farms, we found a high-level diffusion of four efficient colonizing clones (Fig. 6). Their greater ability to disseminate within calves and to persist over time could be attributable to enhanced survival characteristics in the calves’ environment, since *E. coli* is a facultative anaerobic bacterial species which can survive in other habitats than gastrointestinal tracts (72). Once released into the pen, ESBL-producing *E. coli* could be selected on the basis of their ability to implement stress-tolerance mechanisms, which ultimately would result in a subset of clones able to orally re-infect calves. Such mechanisms have been shown to enhance survival in the environment, like a stress-induced mutagenesis mediated by *rpoS* gene (75), a strong ability to produce biofilms (76, 77), and a capacity to grow at temperatures below their optimum (78, 79). Hence, these efficient colonizing clones may have circumvented the potential fitness cost exerted by their *bla*_CTX-M_-carrying plasmid by epistatic interactions between their plasmids and chromosomes. The magnitude of the fitness cost of an antibiotic-resistant carrying plasmid depends on the host genetic background, the plasmid backbone and its gene content, as shown both *in vitro* and *in vivo* (62, 80–82). Studying bacterial host-conjugative plasmid combinations, Silva and colleagues showed that after a period of coevolution, compensatory mutations in antibiotic resistance determinants could arise either on the chromosome or on the plasmid, or both (83). Ultimately, the plasmid-carrying host became fitter than its plasmid-free derivative in 32% of the tested combinations (83). Hence, the antibiotic treatments received by calves may have fostered some ESBL clones at the expense of others, via selection and epistasis compensating *bla*_CTX-M_-carrying plasmid fitness cost. This is in accordance with the successful spread of the clone ‘ST10/466 O8:H32’ (Fig. 6), in which the *mcr-1* gene stably persisted (Fig. 7). Of note, other factors could have also affected the fitness cost of the plasmids, such as the presence of other plasmids in the bacterial host (83, 84), and the animal host species (63).

### Plasmid spread in E. coli clones also drive bla_CTX-M_ dissemination

Another striking result was the ability for some *bla*_CTX-M_-carrying plasmids to spread widely, such as IncI1/ST312 and IncF/F2:A-:B-(Fig. 4, Fig. 5). Spread of *bla*_CTX-M_-carrying plasmid among commensal *E. coli* populations had already been observed in orally inoculated piglets with an *E. coli* strain carrying *bla*_CTX-M-1_ gene (61). It has been shown both *in vitro* and *in vivo* in mice that conjugation frequencies vary across plasmid, donor, and recipient combinations (85). Studies in *Salmonella* have reported the implication of epistatic interactions between the chromosome and plasmids in the spread of the latter in new hosts, involving two-sided active mechanisms (64, 65). Antibiotic treatments may also impact plasmid transfer rates, which is compatible with our observations on *mcr-1*-carrying IncX4 plasmid diffusion in farm A, in which colistin was used (Fig.1, Fig. 7).

Of note, two clones from the CC87, hosting IncI1/ST312 and IncI1/ST3 lineage1 plasmids were detected among the ESBL clones (Fig. S4A). This clonal complex is known to be highly fitted for animal gut environments, enriched in antimicrobial resistance genes and to have a high capacity to acquire and disseminate these genes through conjugation to other members of microbial communities (49). This is in accordance with our findings, as their *bla*_CTX-M_-carrying plasmids were found in several other clones of farm B (Fig. 4). The carriage of *bla*_CTX-M_–carrying plasmids by members of CC87 highlights the implication of clones prone to amplify the dissemination of *bla*_CTX-M_ in veal calf gut microbial communities.

The presence of *bla*_CTX-M_ genes in farm B was mainly driven by plasmid diffusion rather than clonal diffusion. Such gene dynamics observed only in the farm in which no antibiotic treatment was given at start (Fig. 1) may suggest that antibiotic treatment promotes *bla*_CTX-M_ genes persistence by enhancing clone diffusion over plasmid diffusion.

### Multi-farm spreaders were found at low level in each farm

Interestingly, ESBL clones found in several farms were poor colonizers at individual level (Fig. 6), highlighting that even rare ESBL clones in a given ecosystem may encounter an epidemiological success at a macro-level. One of them was of the sequence type ST57, which was also detected in Dutch veal farms (12). In these farms, ST57 was responsible for a bloom of *bla*_CTX-M-14_ spread by an IncF/F2:A-:B-plasmid. The ST57 *E. coli* clone detected in our study also carried *bla*_CTX-M-14_, but on a IncF/F2:A-:B42 plasmid in both farms A and C (Fig. 5). The ST167 was also detected in French farm settings and found prevalent among ESBL-producing *E. coli* isolates (86). It was carrying a *bla*_CTX-M-1_ gene on a IncI1/ST3 plasmid, like in our study. Taken together, these findings show that the local fitness of an ESBL-producing *E. coli* clone cannot be predicted from their dynamics in one farm.

The multi-farm spreader clone ‘ST301/917 O80:H2’ was previously described as an emerging virulent pathotype responsible for hemolytic uremic syndrome and septicemia in humans (50). The serotype O80:H2 carrying the intimin-encoded *eae*-ξ gene was also detected as an emerging pathotype causing diarrhea since 2009 in Belgium calves (87). Although we did not focus our work on pathogenic strains, the presence of this ESBL-producing *E. coli* clone combining pathogenic expression and a phenotype of resistance to last resort antibiotics should be monitored in animals and farmers.

### Limitation of the study

Our study had some limitations. First, it is an observational study conducted in commercial veal farms, which prevent us from drawing conclusions implying causality, specifically regarding the impact of antibiotic treatments administered. Second, samplings started seven days after calves arrived in farms, so that the initial ESBL status of animals remains unknown. Nevertheless, it has been shown that fecal shedding of ESBL-producing *E. coli* clones is high for calves born in dairy farms, from which veal calves originate (88). Moreover, since the vast majority of the 300-350 animals entering a farm are coming from different locations across the French territory, the pool of ESBL-producing *E. coli* clones and plasmids at start is most likely highly diverse. Third, we have sequenced a subset of all the ESBL-producing isolates and obtained a subset of plasmids sequenced after conjugation, limiting our ability to explore more accurately the diversity of ESBL-producing *E. coli* carriage.

In conclusion, we showed that the diffusion of ESBL-encoding genes in calves is a combination of two scenarios encompassing bacterial clone and/or plasmid spread, which highly rely on local features and contexts. In farms where long-term carriage of *bla*_CTX-M_ genes was observed, specific associations between the resistance gene, the plasmid type and the bacterial clone were evidenced. These results argue for complex epistatic interactions between the three parameters, with molecular mechanisms that need to be further understood in the fight against antibiotic resistance.

## DECLARATIONS

## Acknowledgements

We thank Catherine Branger and Luce Landraud for providing strains for qPCR CTX-M, and Caroline Wybraniec for her technical assistance.

## Ethics approval and consent to participate

This study was declared to the CNIL, the French office responsible for protecting personal data, supporting innovation, and preserving individual liberties. No further ethical approval for the use of animals in research was needed since this study did not involve any experimentation on animals (only rectal swabs were sampled) and since we did not collect and register any personal opinion of the participants.

## Data availability

The data for this study have been deposited in the European Nucleotide Archive (ENA) at EMBL-EBI under accession number PRJEB44471 (https://www.ebi.ac.uk/ena/browser/view/PRJEB44471). Raw sequence data are available in the European Nucleotide Archive (EMBL-EMI) (http://www.ebi.ac.uk/ena) under sample accession numbers ERS6288704 to ERS6288746 for isolates sequenced and accession numbers ERS6301044 to ERS6301079 for the transconjugants. The full list and characteristics of these strains are presented in Table S1A and Table S1B along with their genome accession numbers.

## Authors’ contributions

MH, JYM, and ED conceived and designed the study. MH and JYM collected the samples. MH and OC performed the laboratory assays and BC carried out the bioinformatics analyses. MM carried out the statistical analyses of the data and generated the figures. MH and MM wrote the manuscript. MH, JYM, and ED revised and edited the draft. All authors read and approved the final manuscript.

---

**Fig. S1. Distribution of the number of SNPs in bacterial core-genomes according to the delineation of the isolates sequenced in haplogroups and in clones.** Panel A represents the distribution of the number of core-genome Single Nucleotide Polymorphisms (SNPs) between isolates from the same haplogroup or from different haplogroups. An haplogroup is defined as an unique combination of the sequence types (ST) according to the Achtman scheme and the Pasteur Institute scheme, their serotype (O:H) and their *fimH* allele. Each point is the number of SNPs between two core genomes. The number of pairwise comparisons is provided below each group name. Panel B represents the distribution of the number of core-genome SNPs between isolates of the same haplogroup. Panel C represents the pulsed-field gel electrophoresis profile of the four isolates of the haplogroup ‘ST10/2 O101:H9 *fimH54*’ in which we delineate three clones. Panel D represents the distribution of the number of core-genome SNPs between isolates of the same clone. A clone is defined as an unique combination of the phylogroup, the haplogroup, and a number of SNPs between two isolates of the haplogroup inferior to 100 SNP.

**Fig. S2. Distribution of the number of SNPs in plasmid core-genomes and phylogenetic delineation of the IncI1/ST3 plasmids.** For each type of plasmid, panel A represents the distribution of the number of Single Nucleotide Polymorphisms (SNPs) in their core genomes. Plasmids were grouped according to the antibiotic resistance gene they carried (either *bla_CTX-M1_*, *bla_CTX-M14_* or *mcr-1*). Panel B represents the distribution of the number of SNPs in the core genome between the IncI1/ST3 lineages. Panel C represents the genomic content unique to one IncI1/ST3 lineage or shared between lineages (shell and core in the legend). The name of each donor strain is indicated on the x-axis. Plasmids were grouped according to the IncI1/ST3 lineage they belonged to. Shell genome corresponds to coding sequences that were present in more than one IncI1/ST3 lineage, but not all of them. CDS: Coding DNA Sequence, TC: Transconjugant

**Fig. S3A. PFGE profiles of the ESBL-producing *E. coli* isolates in farm A.** Pulsed-field gel electrophoresis (PFGE) was performed on all putative ESBL-producing *E. coli* isolates using the restriction enzyme *Xba*I. Comparison of PFGE profiles was done among isolates to discriminate between ESBL-producing *E. coli* strains circulating in the farm and to select one representative of each PFGE profile for whole genome sequencing. Two isolates were considered to be related to different genotypes if their PFGE profiles differed from at least one band. Isolates having a different status regarding the presence of the *mcr-1* gene compared to their group were additionally selected for genome sequencing. The isolate ID and the sample code (the animal ID followed by the sampling rank) are displayed on the left of the PFGE patterns. On the right, genotypic markers characterized on all isolates, *i.e.* phylogroup, *bla*_CTX-M_ enzyme and presence of the *mcr-1* gene are represented. The clone ID and the *bla*_CTX-M_ gene molecular support are indicated in front of the isolates that were sequenced. Plasmid sequences were obtained from isolates with clone ID in bold after conjugation with a K-12 strain. Isolates of different clones are colored according to the *bla*_CTX-M_ gene molecular support. The isolates 40739, 40742, and 41536 had a smear profile and thus are not depicted in this PFGE figure. Clone identification of the isolates 40739 (‘F ST354/39 O1:H34 *fimH58*’) and 40742 (‘A ST10/466 O8:H32 *fimH23*’) was done by sequencing, while the isolate 41536 was found to be related to the clone ‘A ST10/466 O8:H32 *fimH23*’ through MLVA profile comparisons. COL: Colistin, ESC: Extended-Spectrum Cephalosporins, ‘UG’: unassigned phylogroup.

**Fig. S3B. PFGE profiles of the ESBL-producing *E. coli*isolates in farm B. COL:** Colistin, ESC: Extended-Spectrum Cephalosporins.

**Fig. S3C. PFGE profiles of the ESBL-producing *E. coli* isolates in farm C. COL:** Colistin, ESC: Extended-Spectrum Cephalosporins.

**Fig. S4A. Maximum likelihood phylogenetic tree of ESBL-producing *E. coli* clones.** A representative of each pulsed-field gel electrophoresis (PFGE) profile was selected for whole genome sequencing (WGS). If different *bla*_CTX-M_ genes were detected among a group of isolates having the same PFGE profile, an isolate carrying each of the *bla*_CTX-M_ genes was selected. Isolates carrying the *mcr-1* gene were additionally selected for WGS. The leaves are composed of the isolate ID, then a combination of colored squares representing different O-antigen, H-antigen, and *fimH* gene alleles. The clone was defined by a combination of the phylogroup, ST Achtman/ST Pasteur Institute, serotype, and *fimH* allele gene. The different *bla*_CTX-M_ enzymes detected, the presence of the *mcr-1* gene and the farm in which isolates were sampled are indicated on the right of the tree. The tree was rooted on the *E.coli* ED1a strain, from the B2 phylogroup. It was built using FastTree 2, and is based on the 200,875 SNPs of the 3,003 genes that composed the core genome of this set of 43 genomes. For clarity purpose, the tree presents only 41 isolate IDs because two clones were sequenced in duplicate because of PFGE profile differences (‘A ST10/466 O8:H32 *fimH23*’, ‘A ST301/917 O80:H2 *fimH54*’). ‘SNPs’: Single Nucleotide Polymorphisms, ‘UG’: unassigned phylogroup.

**Fig. S4B. Number of isolates and duration of detection of ESBL-producing *E. coli* clones.** For each farm, the left plot represents the number of times each clone was detected and the right plot represents the days they were isolated. Each color refers to one clone, and the shape represents the *bla*_CTX-M_ gene. The clones are ordered according to their phylogenetic group. The clone was defined by a combination of the phylogroup, ST Achtman/ST Pasteur Institute, serotype, and *fimH* allele gene.

**Fig. S4C. Resistome of ESBL-producing *E. coli* clones.** Heatmap of the antibiotic resistance genes detected in the ESBL-producing *E. coli* clones using the ResFinder database. Resistance genes are indicated on the x-axis and are grouped per class of antibiotics. *E. coli* clones are indicated on the y-axis and are grouped according to the farm in which they were isolated. The clone was defined by a combination of the phylogroup, ST Achtman/ST Pasteur Institute, serotype, and *fimH* allele gene. Resistance genes were considered to be carried by a clone if it was found in at least one sequenced genome of this clone. The number of isolates per clone in each farm is displayed on the right of the plot. ‘UG’: unassigned phylogroup.

**Fig. S5A. Distribution of extraintestinal pathogenic virulence genes detected in ESBL-producing *E. coli* clones.** Heatmap of the virulence genes associated with extraintestinal pathogenic *E. coli*, using the VFDB and VirulenceFinder databases. Virulence genes are indicated on the x-axis and are grouped per function encoded. ESBL-producing *E. coli* clones are indicated on the y-axis and are grouped according to the farm in which they were isolated. The clone was defined by a combination of the phylogroup, ST Achtman/ST Pasteur Institute, serotype, and *fimH* allele gene. The number of extraintestinal pathogenic virulence genes detected in each clone on the right.

**Fig. S5B. Distribution of intestinal pathogenic virulence genes detected in ESBL-producing *E. coli* clones.** Heatmap of the virulence genes associated with intestinal pathogenic *E. coli*, using the VFDB and VirulenceFinder databases. Virulence genes are indicated on the x-axis and are grouped per function encoded. ESBL-producing *E. coli* clones are indicated on the y-axis and are grouped according to the farm in which they were isolated. The clone ID is a combination of the phylogroup, ST Achtman/ST Pasteur Institute, serotype, and *fimH* allele gene. The number of intestinal pathogenic virulence genes detected in each clone is displayed on the right.

**Fig. S5C. Distribution of pathogenic virulence genes associated with both extraintestinal and intestinal pathogenicity, and bacteriocins detected in ESBL-producing *E. coli* clones.** Heatmap of the virulence genes associated with both extraintestinal and intestinal pathogenic *E. coli*, using the VFDB and VirulenceFinder databases and of the bacteriocins detected in the isolates sequenced. Virulence genes are indicated on the x-axis and are grouped per function encoded. ESBL-producing *E. coli* clones are indicated on the y-axis and are grouped according to the farm in which they were isolated. The clone ID is a combination of the phylogroup, ST Achtman/ST Pasteur Institute, serotype, and *fimH* allele gene. The number of virulence genes detected in each clone is displayed on the right.

**Fig. S6A. Linear map of the chromosomal environment of *bla*_CTX-M-15_ gene and *mcr-1* gene.** The chromosomal environment of the *bla*_CTX-M-15_ gene found in the clone ‘A ST10/2 O101:H9 *fimH54* clone2’ isolated in farms B and C (isolate IDs 40772 and 41054, respectively) is represented on the top of the figure. The chromosomal environments of the *mcr-1* gene in clones ‘F ST354/39 O1:H34 fimH58’ in farm A (isolate ID 40808) and ‘G ST117/48 O24:H4 fimH189’ in farm C (isolate ID 41301), are represented on the middle and the bottom of the figure, respectively. The antibiotic resistance genes are indicated by red arrows. Open reading frames are shown as arrows indicating the direction of transcription (dark blue, plasmid transfer; red, resistance; pink, mobile elements). The name of genes is indicated within the arrows. The genome references against which contigs containing the genes were blasted are indicated below the maps.

**Fig. S6B. Circular maps of prevalent plasmids carrying *bla*_CTX-M_ genes and *mcr-1* gene.** The genome of transconjugants (TC) carrying plasmid-borne ESBL-encoding genes or *mcr-1* gene were sequenced to characterize the molecular environment of these antibiotic resistance genes. Plasmids were identified using the PlasmidFinder database and reference genomes were found by blasting contigs against MaGe or Refseq databases. The map and size of plasmids IncI1/ST312, IncF/F2:A-:B-, and IncX4 are represented on the left, the middle and the right of the plot, respectively. Open reading frames are shown as boxes colored according to the function of their product. ESBL-encoding genes (*bla*_CTX-M-1_ and *bla*_CTX-M-14_ genes) and *mcr-1* gene are indicated in red on the map. Single Nucleotide Polymorphisms (SNPs) detected in the plasmidic contigs of the TC are represented by black lines. The donor strain ID of each TC is given below the figure, and the farm in which it was isolated is provided between brackets.

**Fig. S6C. Circular maps of plasmids carrying *bla*_CTX-M_ genes.** The genome of transconjugants (TC) carrying plasmid-borne ESBL-encoding genes or *mcr-1* gene were sequenced to characterize the molecular environment of these antibiotic resistance genes. Plasmids were identified using the PlasmidFinder database and reference genomes were found by blasting contigs against MaGe or Refseq databases. The map and size of plasmids IncI1/ST3, IncF/F59:A-:B-, and IncK are represented on the left, the middle and the right of the plot, respectively. Open reading frames are shown as boxes colored according to the function of their product. ESBL-encoding genes (*bla*_CTX-M-1_ and *bla*_CTX-M-14_ genes) and *mcr-1* gene are indicated in red on the map. Single Nucleotide Polymorphisms (SNPs) and deletions detected in the plasmidic contigs of the TC are represented by black and white lines, respectively. The donor strain ID of each TC is given below the figure, and the farm in which it was isolated is provided between brackets. For IncI1/ST3 plasmid, the lineage of each plasmid sequenced is indicated below the figure.

**Table S1A. Summary of the origin and molecular characteristics of the ESBL-producing E. coli isolates and plasmids.** Pulsed-field gel electrophoresis (PFGE) was performed on all putative ESBL-producing *E. coli* isolates using the restriction enzyme *Xba*I. Comparison of PFGE profiles was done among isolates to discriminate between ESBL-producing *E. coli* strains circulating in the farm and to select one representative of each PFGE profile for whole genome sequencing. Two isolates were considered to be related to different genotypes if their PFGE profiles differed from at least one band. Isolates having a different status regarding the presence of the *mcr-1* gene compared to their group were additionally selected for genome sequencing. The clone ID and the sample code (the animal ID followed by the sampling rank) are also presented in the table.

**Table S1B. Summary of the origin and molecular characteristics of the plasmids carrying bla_CTX-M_ genes and mcr-1 gene.** The genome of transconjugants (TC) carrying plasmid-borne ESBL-encoding genes or *mcr-1* gene were sequenced to characterize the molecular environment of these antibiotic resistance genes. Plasmids were identified using the PlasmidFinder database and reference genomes were found by blasting contigs against MaGe or Refseq databases.

**Table S2. Posthoc Dunn’s test results, comparison of the number of copies of *bla*_CTX-M_ genes (log_10_(copies */* g)) estimated by quantitative PCR between the samples grouped according to the number of colonies on ESBL-producing *Enterobacteriaceae* selective medium.** Quantification of *bla*_CTX-M_ group 1 and *bla*_CTX-M_ group 9 gene copies was performed by qPCR on samples collected 7 days and 21 days after arrival in farms, and one by month for each calf.

**Table S3A.** Phenotypic prevalence of antibiotic co-resistances in the ESBL-producing *E. coli* isolates (n= 173). Antibiotics are sorted in descending order according to their frequency and classes.

**Table S3B. Quinolone resistance mutations detected in the ESBL-producing *E. coli* isolates.** Quinolone resistance point mutations were searched using the website “Center for Genomic Epidemiology” with standard parameters.

