## Supplementary Figure S4 for "Interplay between bacterial clone and plasmid in the spread of antibiotic resistance genes in the gut: lessons from a temporal study in veal calves"

**Fig. S4A. Maximum likelihood phylogenetic tree of ESBL-producing *E. coli* clones.** A representative of each pulsed-field gel electrophoresis (PFGE) profile was selected for whole genome sequencing (WGS). If different $bla_{\text{CTX-M}}$ genes were detected among a group of isolates having the same PFGE profile, an isolate carrying each of the $bla_{\text{CTX-M}}$ genes was selected. Isolates carrying the *mcr-1* gene were additionally selected for WGS. The leaves are composed of the isolate ID, then a combination of colored squares representing different O-antigen, H-antigen, and *fimH* gene alleles. The clone was defined by a combination of the phylogroup, ST Achtman/ST Pasteur Institute, serotype, and *fimH* allele gene. The different $bla_{\text{CTX-M}}$ enzymes detected, the presence of the *mcr-1* gene and the farm in which isolates were sampled are indicated on the right of the tree. The tree was rooted on the *E.coli* ED1a strain, from the B2 phylogroup. It was built using FastTree 2, and is based on the 200,875 SNPs of the 3,003 genes that composed the core genome of this set of 43 genomes. For clarity purpose, the tree presents only 41 isolate IDs because two clones were sequenced in duplicate because of PFGE profile differences (‘A ST10/466 O8:H32 *fimH23*’, ‘A ST301/917 O80:H2 *fimH54*’). ‘SNPs’: Single Nucleotide Polymorphisms, ‘UG’: unassigned phylogroup.

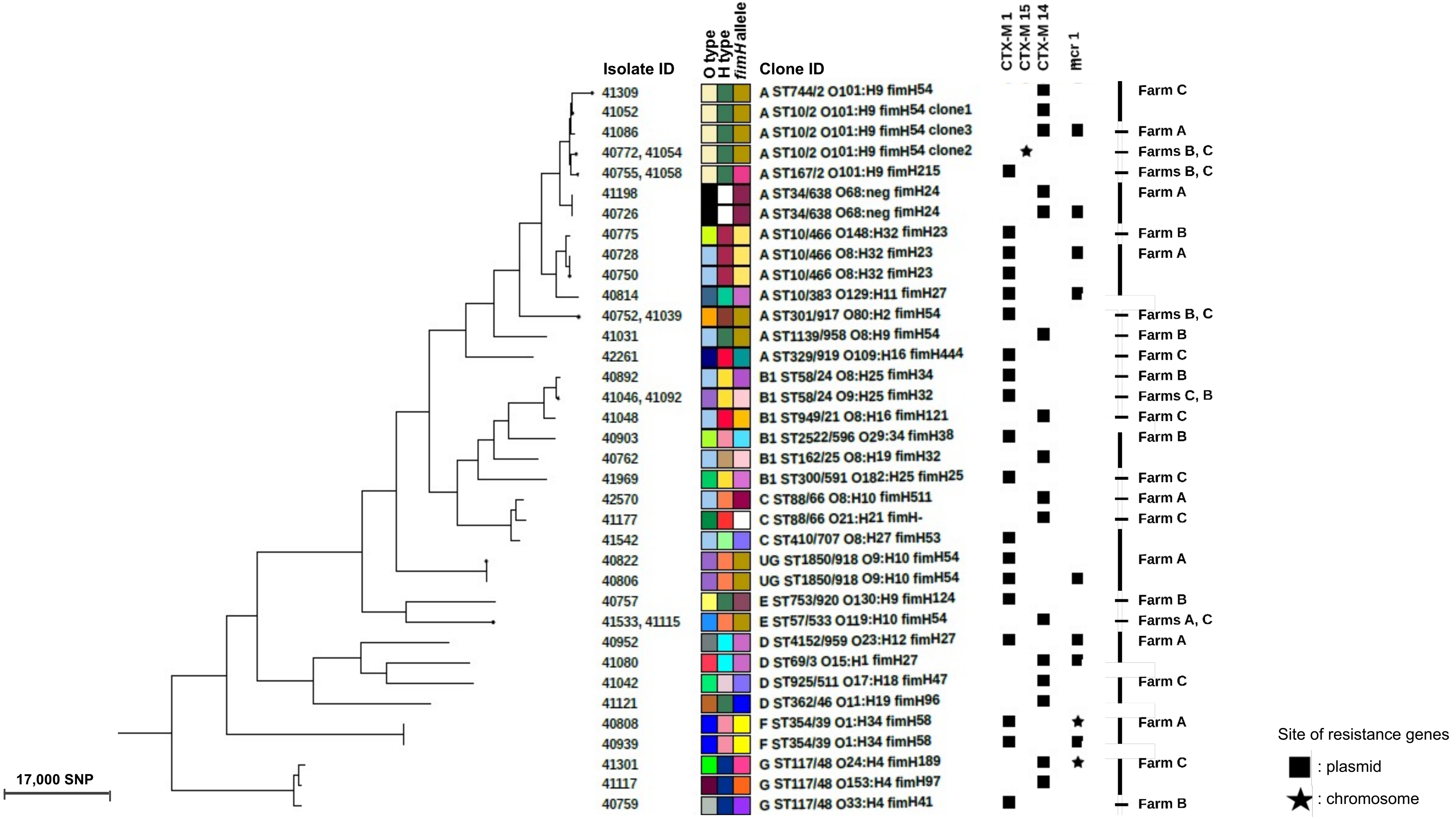

**Fig. S4B. Number of isolates and duration of detection of ESBL-producing *E. coli* clones.** For each farm, the left plot represents the number of times each clone was detected and the right plot represents the days they were isolated. Each color refers to one clone, and the shape represents the *bla*$_{CTX-M}$ gene. The clones are ordered according to their phylogenetic group. The clone was defined by a combination of the phylogroup, ST Achtman/ST Pasteur Institute, serotype, and *fimH* allele gene.

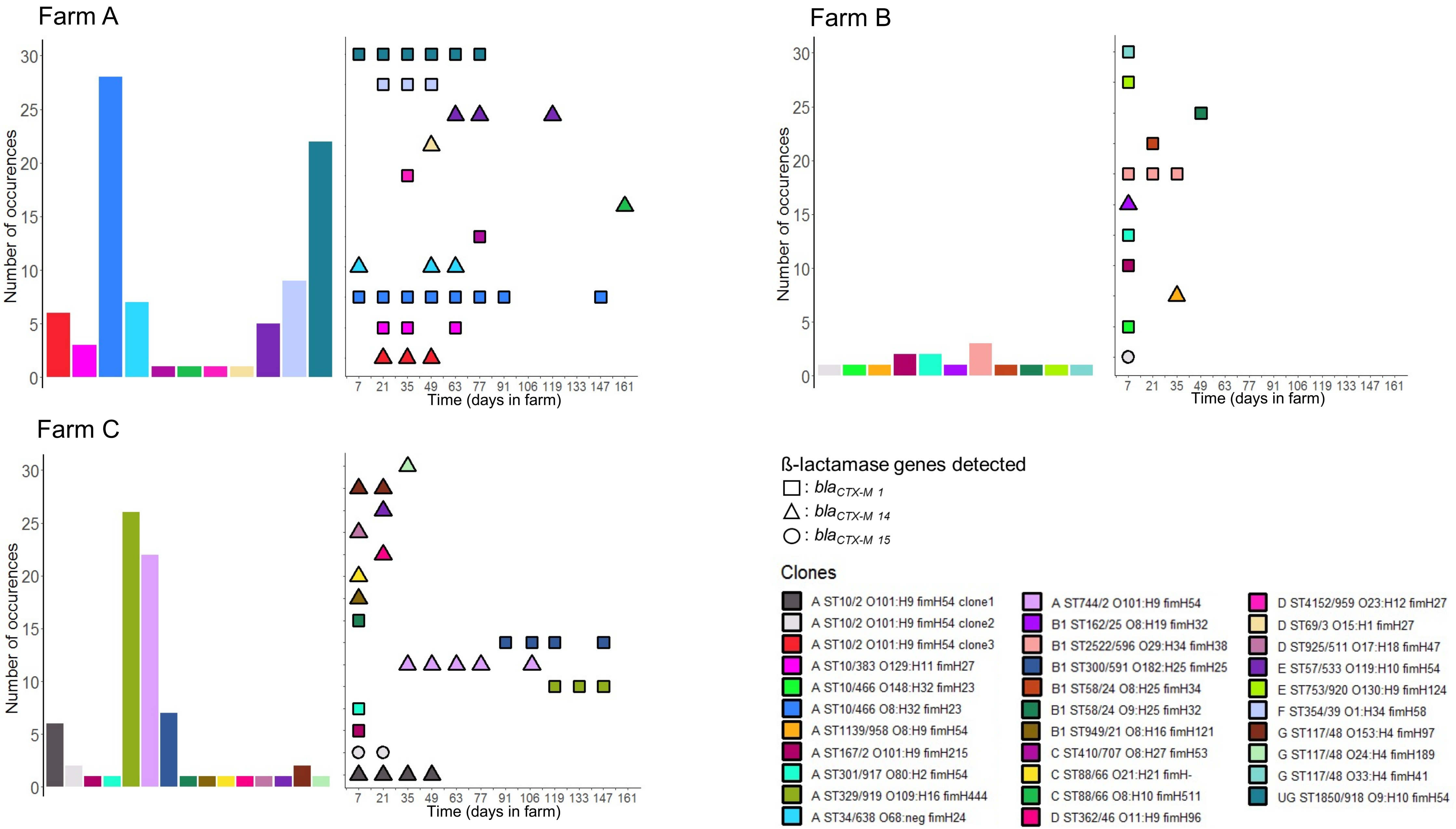

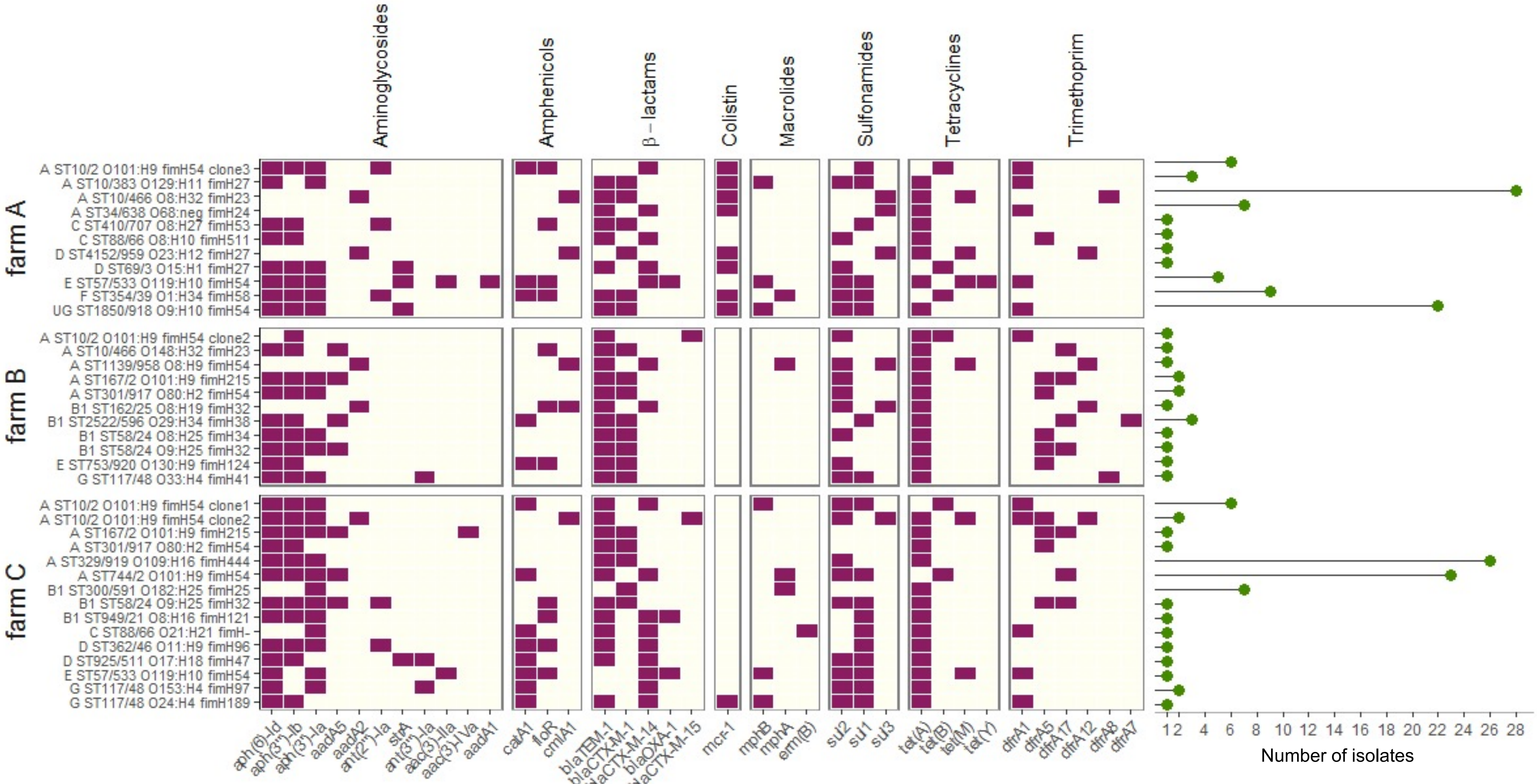


**Fig. S4C. Resistome of ESBL-producing *E. coli* clones.** Heatmap of the antibiotic resistance genes detected in the ESBL-producing *E. coli* clones using the ResFinder database. Resistance genes are indicated on the x-axis and are grouped per class of antibiotics. E. coli clones are indicated on the y-axis and are grouped according to the farm in which they were isolated. The clone was defined by a combination of the phylogroup, ST Achtman/ST Pasteur Institute, serotype, and fimH allele gene. Resistance genes were considered to be carried by a clone if it was found in at least one sequenced genome of this clone. The number of isolates per clone in each farm is displayed on the right of the plot. 'UG': unassigned phylogroup.
