## Supplementary Figures S1, S3A, S3C for "Interplay between bacterial clone and plasmid in the spread of antibiotic resistance genes in the gut: lessons from a temporal study in veal calves"

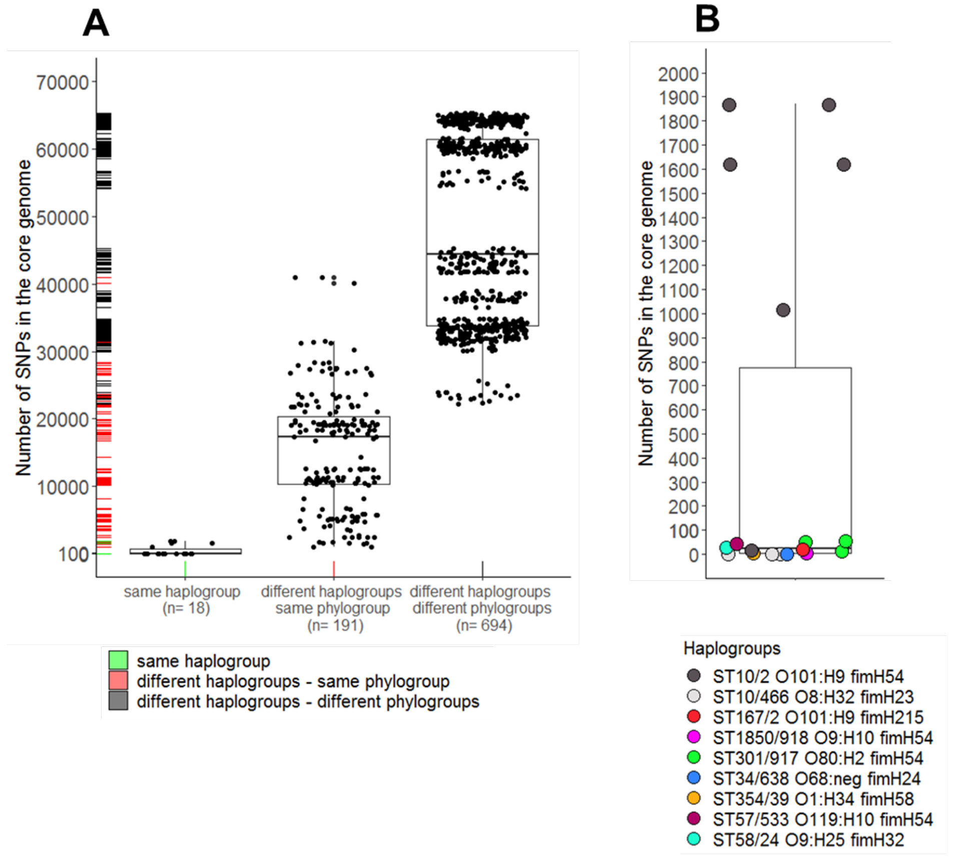


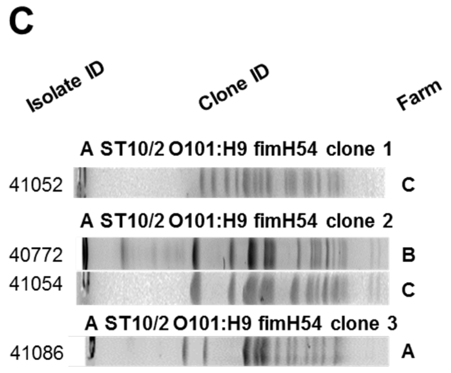


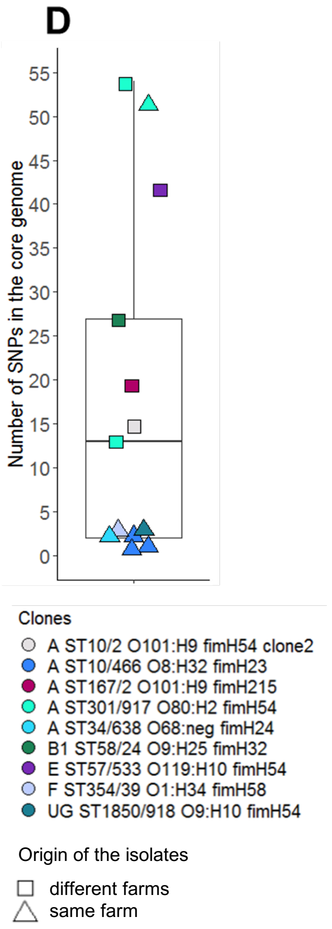


**Fig. S1. Distribution of the number of SNPs in bacterial core-genomes according to the delineation of the isolates sequenced in haplogroups and in clones.** Panel A represents the distribution of the number of core-genome Single Nucleotide Polymorphisms (SNPs) between isolates from the same haplogroup or from different haplogroups. An haplogroup is defined as an unique combination of the sequence types (ST) according to the Achtman scheme and the Pasteur Institute scheme, their serotype (O:H) and their *fimH* allele. Each point is the number of SNPs between two core genomes. The number of pairwise comparisons is provided below each group name. Panel B represents the distribution of the number of core-genome SNPs between isolates of the same haplogroup. Panel C represents the pulsed-field gel electrophoresis profile of the four isolates of the haplogroup 'ST10/2 O101:H9 *fimH54*' in which we delineate three clones. Panel D represents the distribution of the number of core-genome SNPs between isolates of the same clone. A clone is defined as an unique combination of the phylogroup, the haplogroup, and a number of SNPs between two isolates of the haplogroup inferior to 100 SNP.

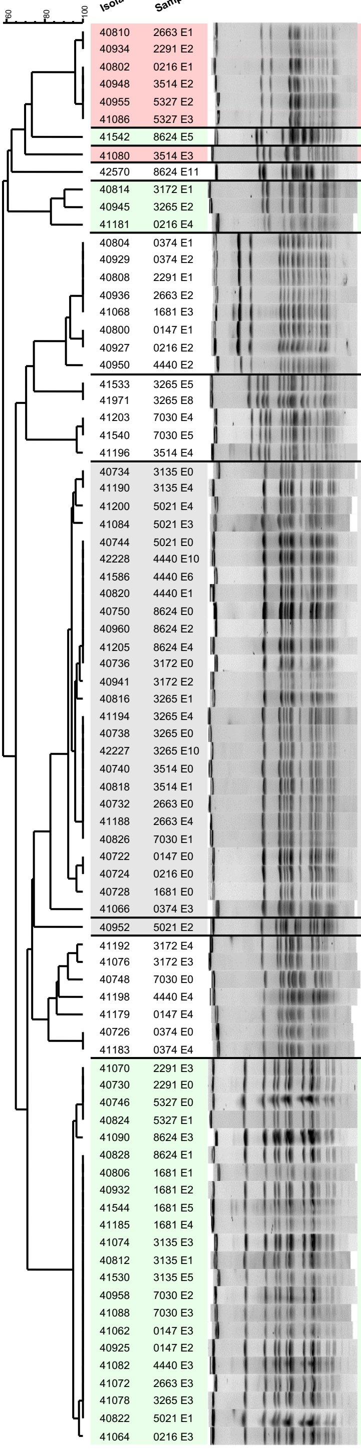


| Isolate ID | Sample code | Phylogroup | Clone ID | ESC-resistance enzyme | COL-resistance enzyme | $bla_{CTX-M}$ gene location |
|---|---|---|---|---|---|---|
| 40810 | 2663 E1 | A | | CTX-M-14 | MCR-1 | |
| 40934 | 2291 E2 | A | | CTX-M-14 | MCR-1 | |
| 40802 | 0216 E1 | A | | CTX-M-14 | MCR-1 | |
| 40948 | 3514 E2 | A | | CTX-M-14 | MCR-1 | |
| 40955 | 5327 E2 | A | | CTX-M-14 | MCR-1 | |
| 41086 | 5327 E3 | A | **A ST10/2 O101:H9 *fimH54* clone3** | CTX-M-14 | MCR-1 | IncF/F2:A-:B- |
| 41542 | 8624 E5 | C | **C ST410/707 O8:H27 *fimH53*** | CTX-M-1 | - | IncI1/ST312 |
| 41080 | 3514 E3 | D | **D ST69/3 O15:H1 *fimH27*** | CTX-M-14 | MCR-1 | IncF/F2:A-:B- |
| 42570 | 8624 E11 | C | **C ST88/66 O8:H10 *fimH511*** | CTX-M-14 | - | IncK |
| 40814 | 3172 E1 | A | **A ST10/383 O129:H11 *fimH27*** | CTX-M-1 | MCR-1 | IncI1/ST312 |
| 40945 | 3265 E2 | A | | CTX-M-1 | - | |
| 41181 | 0216 E4 | A | | CTX-M-1 | - | |
| 40804 | 0374 E1 | F | | CTX-M-1 | MCR-1 | |
| 40929 | 0374 E2 | F | | CTX-M-1 | MCR-1 | |
| 40808 | 2291 E1 | F | F ST354/39 O1:H34 *fimH58* | CTX-M-1 | MCR-1 | unknown |
| 40936 | 2663 E2 | F | | CTX-M-1 | MCR-1 | |
| 41068 | 1681 E3 | F | | CTX-M-1 | MCR-1 | |
| 40800 | 0147 E1 | F | | CTX-M-1 | MCR-1 | |
| 40927 | 0216 E2 | F | | CTX-M-1 | MCR-1 | |
| 40950 | 4440 E2 | F | | CTX-M-1 | MCR-1 | |
| 41533 | 3265 E5 | E | **E ST57/533 O119:H10 *fimH54*** | CTX-M-14 | - | IncF/F2:A-:B42 |
| 41971 | 3265 E8 | E | | CTX-M-14 | - | |
| 41203 | 7030 E4 | E | | CTX-M-14 | - | |
| 41540 | 7030 E5 | E | | CTX-M-14 | - | |
| 41196 | 3514 E4 | E | | CTX-M-14 | - | |
| 40734 | 3135 E0 | A | | CTX-M-1 | MCR-1 | |
| 41190 | 3135 E4 | A | | CTX-M-1 | MCR-1 | |
| 41200 | 5021 E4 | A | | CTX-M-1 | MCR-1 | |
| 41084 | 5021 E3 | A | | CTX-M-1 | MCR-1 | |
| 40744 | 5021 E0 | A | | CTX-M-1 | MCR-1 | |
| 42228 | 4440 E10 | A | | CTX-M-1 | MCR-1 | |
| 41586 | 4440 E6 | A | | CTX-M-1 | MCR-1 | |
| 40820 | 4440 E1 | A | | CTX-M-1 | MCR-1 | |
| 40750 | 8624 E0 | A | **A ST10/466 O8:H32 *fimH23*** | CTX-M-1 | - | IncF/F59:A-:B- |
| 40960 | 8624 E2 | A | | CTX-M-1 | MCR-1 | |
| 41205 | 8624 E4 | A | | CTX-M-1 | MCR-1 | |
| 40736 | 3172 E0 | A | | CTX-M-1 | MCR-1 | |
| 40941 | 3172 E2 | A | | CTX-M-1 | MCR-1 | |
| 40816 | 3265 E1 | A | | CTX-M-1 | MCR-1 | |
| 41194 | 3265 E4 | A | | CTX-M-1 | MCR-1 | |
| 40738 | 3265 E0 | A | | CTX-M-1 | MCR-1 | |
| 42227 | 3265 E10 | A | | CTX-M-1 | MCR-1 | |
| 40740 | 3514 E0 | A | | CTX-M-1 | MCR-1 | |
| 40818 | 3514 E1 | A | | CTX-M-1 | MCR-1 | |
| 40732 | 2663 E0 | A | | CTX-M-1 | MCR-1 | |
| 41188 | 2663 E4 | A | | CTX-M-1 | MCR-1 | |
| 40826 | 7030 E1 | A | | CTX-M-1 | MCR-1 | |
| 40722 | 0147 E0 | A | | CTX-M-1 | MCR-1 | |
| 40724 | 0216 E0 | A | | CTX-M-1 | MCR-1 | |
| 40728 | 1681 E0 | A | **A ST10/466 O8:H32 *fimH23*** | CTX-M-1 | MCR-1 | IncF/F59:A-:B- |
| 41066 | 0374 E3 | A | | CTX-M-1 | MCR-1 | |
| 40952 | 5021 E2 | D | D ST4152/959 O23:H12 *fimH27* | CTX-M-1 | MCR-1 | IncF/F59:A-:B- |
| 41192 | 3172 E4 | A | | CTX-M-14 | MCR-1 | |
| 41076 | 3172 E3 | A | | CTX-M-14 | MCR-1 | |
| 40748 | 7030 E0 | A | | CTX-M-14 | MCR-1 | |
| 41198 | 4440 E4 | A | A ST34/638 O68:neg *fimH24* | CTX-M-14 | - | |
| 41179 | 0147 E4 | A | | CTX-M-14 | - | |
| 40726 | 0374 E0 | A | **A ST34/638 O68:neg *fimH24*** | CTX-M-14 | MCR-1 | IncF/F2:A-:B49 |
| 41183 | 0374 E4 | A | | CTX-M-14 | MCR-1 | |
| 41070 | 2291 E3 | UG | | CTX-M-1 | - | |
| 40730 | 2291 E0 | UG | | CTX-M-1 | - | |
| 40746 | 5327 E0 | UG | | CTX-M-1 | MCR-1 | |
| 40824 | 5327 E1 | UG | | CTX-M-1 | - | |
| 41090 | 8624 E3 | UG | | CTX-M-1 | - | |
| 40828 | 8624 E1 | UG | | CTX-M-1 | - | |
| 40806 | 1681 E1 | UG | **UG ST1850/918 O9:H10 *fimH54*** | CTX-M-1 | MCR-1 | IncI1/ST312 |
| 40932 | 1681 E2 | UG | | CTX-M-1 | - | |
| 41544 | 1681 E5 | UG | | CTX-M-1 | - | |
| 41185 | 1681 E4 | UG | | CTX-M-1 | - | |
| 41074 | 3135 E3 | UG | | CTX-M-1 | - | |
| 40812 | 3135 E1 | UG | | CTX-M-1 | - | |
| 41530 | 3135 E5 | UG | | CTX-M-1 | - | |
| 40958 | 7030 E2 | UG | | CTX-M-1 | - | |
| 41088 | 7030 E3 | UG | | CTX-M-1 | - | |
| 41062 | 0147 E3 | UG | | CTX-M-1 | - | |
| 40925 | 0147 E2 | UG | | CTX-M-1 | - | |
| 41082 | 4440 E3 | UG | | CTX-M-1 | - | |
| 41072 | 2663 E3 | UG | | CTX-M-1 | - | |
| 41078 | 3265 E3 | UG | | CTX-M-1 | - | |
| 40822 | 5021 E1 | UG | **UG ST1850/918 O9:H10 *fimH54*** | CTX-M-1 | - | IncI1/ST312 |
| 41064 | 0216 E3 | UG | | CTX-M-1 | - | |

**Fig. S3A. PFGE profiles of the ESBL-producing *E. coli* isolates in farm A.** Pulsed-field gel electrophoresis (PFGE) was performed on all putative ESBL-producing *E. coli* isolates using the restriction enzyme *Xba*I. Comparison of PFGE profiles was done among isolates to discriminate between ESBL-producing *E. coli* strains circulating in the farm and to select one representative of each PFGE profile for whole genome sequencing. Two isolates were considered to be related to different genotypes if their PFGE profiles differed from at least one band. Isolates having a different status regarding the presence of the *mcr-1* gene compared to their group were additionally selected for genome sequencing. The isolate ID and the sample code (the animal ID followed by the sampling rank) are displayed on the left of the PFGE patterns. On the right, genotypic markers characterized on all isolates, *i.e.* phylogroup, $bla_{CTX-M}$ enzyme and presence of the *mcr-1* gene are represented. The clone ID and the $bla_{CTX-M}$ gene molecular support are indicated in front of the isolates that were sequenced. Plasmid sequences were obtained from isolates with clone ID in bold after conjugation with a K-12 strain. Isolates of different clones are colored according to the $bla_{CTX-M}$ gene molecular support. The isolates 40739, 40742, and 41536 had a smear profile and thus are not depicted in this PFGE figure. Clone identification of the isolates 40739 ('F ST354/39 O1:H34 *fimH58*') and 40742 ('A ST10/466 O8:H32 *fimH23*') was done by sequencing, while the isolate 41536 was found to be related to the clone 'A ST10/466 O8:H32 *fimH23*' through MLVA profile comparisons. COL: Colistin, ESC: Extended-Spectrum Cephalosporins, 'UG': unassigned phylogroup.

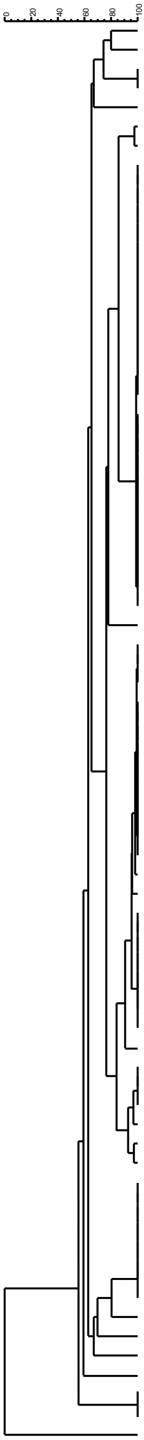


| Isolate ID | Sample code | Phylogroup | Clone ID | ESC-resistance enzyme | COL-resistance enzyme | $bla_{CTX-M}$ gene location |
|---|---|---|---|---|---|---|
| 41115 | 1232 E1 | E | **E ST57/533 O119:H10 *fimH54*** | CTX-M-14 | - | IncF/F2:A-:B42 |
| 41301 | 2235 E2 | G | **G ST117/48 O24:H4 *fimH189*** | CTX-M-14 | MCR-1 | IncF/F2:A-:B- |
| 41044 | 2235 E0 | G | | CTX-M-14 | - | |
| 41117 | 2235 E1 | G | **G ST117/48 O153:H4 *fimH97*** | CTX-M-14 | - | IncF/F2:A-:B- |
| 41177 | 0424 E0 | C | **C ST88/66 O21:H21 *fimH-*** | CTX-M-14 | - | IncI1/ST80 |
| 42608 | 5614 E9 | A | | CTX-M-1 | - | |
| 42631 | 5614 E10 | A | | CTX-M-1 | - | |
| 42266 | 5614 E8 | A | | CTX-M-1 | - | |
| 42611 | 7091 E9 | A | | CTX-M-1 | - | |
| 42269 | 7091 E8 | A | | CTX-M-1 | - | |
| 42263 | 2391 E8 | A | | CTX-M-1 | - | |
| 42604 | 2391 E9 | A | | CTX-M-1 | - | |
| 42267 | 6230 E8 | A | | CTX-M-1 | - | |
| 42609 | 6230 E9 | A | | CTX-M-1 | - | |
| 42268 | 6567 E8 | A | | CTX-M-1 | - | |
| 42610 | 6567 E9 | A | | CTX-M-1 | - | |
| 42262 | 2235 E8 | A | | CTX-M-1 | - | |
| 42603 | 2235 E9 | A | | CTX-M-1 | - | |
| 42258 | 0191 E8 | A | | CTX-M-1 | - | |
| 42600 | 0191 E9 | A | | CTX-M-1 | - | |
| 42601 | 0196 E9 | A | | CTX-M-1 | - | |
| 42259 | 0196 E8 | A | | CTX-M-1 | - | |
| 42629 | 0196 E10 | A | | CTX-M-1 | - | |
| 42265 | 3010 E8 | A | | CTX-M-1 | - | |
| 42606 | 3010 E9 | A | | CTX-M-1 | - | |
| 42260 | 0424 E8 | A | | CTX-M-1 | - | |
| 42602 | 0424 E9 | A | | CTX-M-1 | - | |
| 42261 | 1232 E8 | A | **A ST329/919 O109:H16 *fimH444*** | CTX-M-1 | - | IncI1/ST3 lineage 2 |
| 42270 | 9826 E8 | A | | CTX-M-1 | - | |
| 42607 | 5071 E9 | A | | CTX-M-1 | - | |
| 42605 | 2827 E9 | A | | CTX-M-1 | - | |
| 41048 | 2827 E0 | B1 | **B1 ST949/21 O8:H16 *fimH121*** | CTX-M-14 | - | IncF/F2:A-:B- |
| 41313 | 6230 E2 | A | | CTX-M-14 | - | |
| 41619 | 6230 E4 | A | | CTX-M-14 | - | |
| 41315 | 6567 E2 | A | | CTX-M-14 | - | |
| 41571 | 6567 E3 | A | | CTX-M-14 | - | |
| 41621 | 6567 E4 | A | | CTX-M-14 | - | |
| 41297 | 0196 E2 | A | | CTX-M-14 | - | |
| 41608 | 0196 E4 | A | | CTX-M-14 | - | |
| 41307 | 3010 E2 | A | | CTX-M-14 | - | |
| 41614 | 3010 E4 | A | | CTX-M-14 | - | |
| 41616 | 5071 E4 | A | | CTX-M-14 | - | |
| 42092 | 5071 E7 | A | | CTX-M-14 | - | |
| 41309 | 5071 E2 | A | | CTX-M-14 | - | |
| 41319 | 9826 E2 | A | | CTX-M-14 | - | |
| 41624 | 9826 E4 | A | | CTX-M-14 | - | |
| 41305 | 2827 E2 | A | | CTX-M-14 | - | |
| 41778 | 6567 E5 | A | | CTX-M-14 | - | |
| 41321 | 0424 E2 | A | | CTX-M-14 | - | |
| 41311 | 5614 E2 | A | | CTX-M-14 | - | |
| 41317 | 7091 E2 | A | | CTX-M-14 | - | |
| 41293 | 0191 E2 | A | | CTX-M-14 | - | |
| 42350 | 2235 E4 | A | | CTX-M-14 | - | |
| 41299 | 1232 E2 | A | **A ST744/2 O101:H9 *fimH54*** | CTX-M-14 | - | IncK |
| 41050 | 3010 E0 | A | | CTX-M-14 | - | |
| 41052 | 5071 E0 | A | **A ST10/2 O101:H9 *fimH54* clone1** | CTX-M-14 | - | IncF/F2:A-:B- |
| 41112 | 0196 E1 | A | | CTX-M-14 | - | |
| 41056 | 6230 E0 | A | | CTX-M-14 | - | |
| 41303 | 2391 E2 | A | | CTX-M-14 | - | |
| 42349 | 2235 E3 | A | | CTX-M-14 | - | |
| 41969 | 1232 E6 | B1 | B1 ST300/591 O182:H25 *fimH25* | CTX-M-1 | - | unknown |
| 42093 | 5614 E7 | B1 | | CTX-M-1 | - | |
| 42094 | 6230 E7 | B1 | | CTX-M-1 | - | |
| 42095 | 6567 E7 | B1 | | CTX-M-1 | - | |
| 42264 | 2827 E8 | B1 | | CTX-M-1 | - | |
| 42628 | 0191 E10 | B1 | | CTX-M-1 | - | |
| 42630 | 0424 E10 | B1 | | CTX-M-1 | - | |
| 41039 | 0196 E0 | A | A ST301/917 O80:H2 *fimH54* | CTX-M-1 | - | IncI1/ST3 (lineage unknown) |
| 41046 | 2391 E0 | B1 | **B1 ST58/24 O9:H25 *fimH32*** | CTX-M-1 | - | IncI1/ST3 lineage 3 |
| 41121 | 2827 E1 | D | **D ST362/46 O11:H19 *fimH96*** | CTX-M-14 | - | IncF/F2:A-:B- |
| 41058 | 6567 E0 | A | A ST167/2 O101:H9 *fimH215* | CTX-M-1 | - | unknown |
| 41054 | 5614 E0 | A | A ST10/2 O101:H9 *fimH54* clone2 | CTX-M-15 | - | chromosomic |
| 41125 | 5614 E1 | A | | CTX-M-15 | - | |
| 41042 | 1232 E0 | D | **D ST925/511 O17:H18 *fimH47*** | CTX-M-14 | - | IncF/F2:A-:B- |

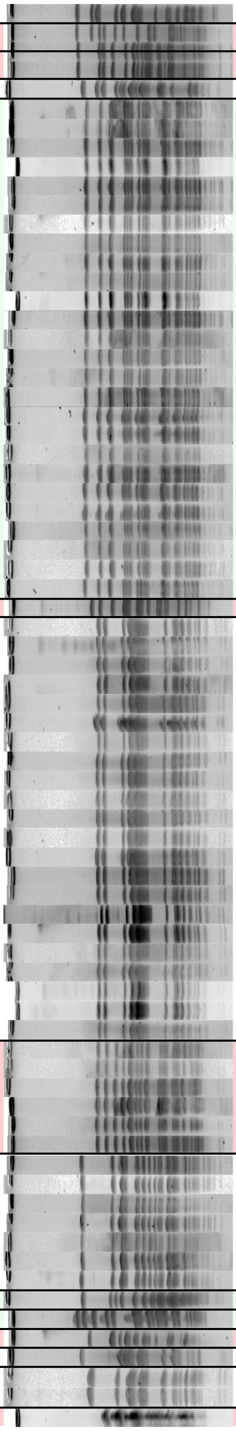

**Fig. S3C. PFGE profiles of the ESBL-producing *E. coli* isolates in farm C.** Pulsed-field gel electrophoresis (PFGE) was performed on all putative ESBL-producing *E. coli* isolates using the restriction enzyme *Xba*I. Comparison of PFGE profiles was done among isolates to discriminate between ESBL-producing *E. coli* strains circulating in the farm and to select one representative of each PFGE profile for genome sequencing. Two isolates were considered to be related to different genotypes if their PFGE profiles differed from at least one band. Isolates having a different status regarding the presence of the *mcr-1* gene compared to their group were additionally selected for genome sequencing. The isolate ID and the sample code (the animal ID followed by the sampling rank) are displayed on the left of the PFGE patterns. On the right, genotypic markers characterized on all isolates, *i.e.* phylogroup, $bla_{CTX-M}$ enzyme and presence of the *mcr-1* gene are represented. The clone ID and the $bla_{CTX-M}$ gene molecular support are indicated in front of the isolates that were sequenced. Plasmid sequences were obtained from isolates with clone ID in bold after conjugation with a K-12 strain. Isolates of different clones are colored according to the $bla_{CTX-M}$ gene molecular support. COL: Colistin, ESC: Extended-Spectrum Cephalosporins.
