## Supplementary Figures S2 S3B for "Interplay between bacterial clone and plasmid in the spread of antibiotic resistance genes in the gut: lessons from a temporal study in veal calves"

A

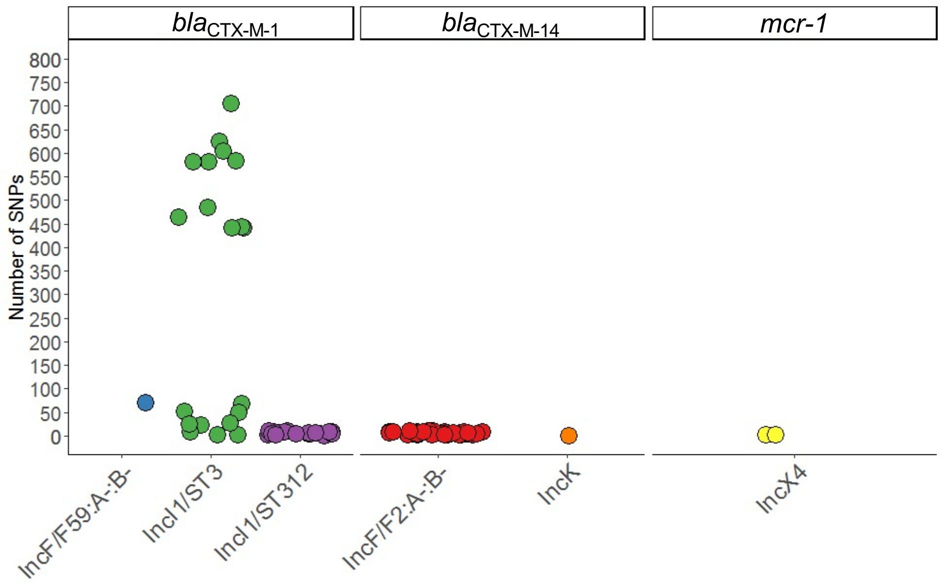


**Fig. S2. Distribution of the number of SNPs in plasmid core-genomes and phylogenetic delineation of the IncI1/ST3 plasmids.** For each type of plasmid, panel A represents the distribution of the number of Single Nucleotide Polymorphisms (SNPs) in their core genomes. Plasmids were grouped according to the antibiotic resistance gene they carried (either *bla*$_{CTX\text{-}M1}$, *bla*$_{CTX\text{-}M14}$ or *mcr-1*). Panel B represents the distribution of the number of SNPs in the core genome between the IncI1/ST3 lineages. Panel C represents the genomic content unique to one IncI1/ST3 lineage or shared between lineages (shell and core in the legend). The name of each donor strain is indicated on the x-axis. Plasmids were grouped according to the IncI1/ST3 lineage they belonged to. Shell genome corresponds to coding sequences that were present in more than one IncI1/ST3 lineage, but not all of them. CDS: Coding DNA Sequence, TC: Transconjugant

B

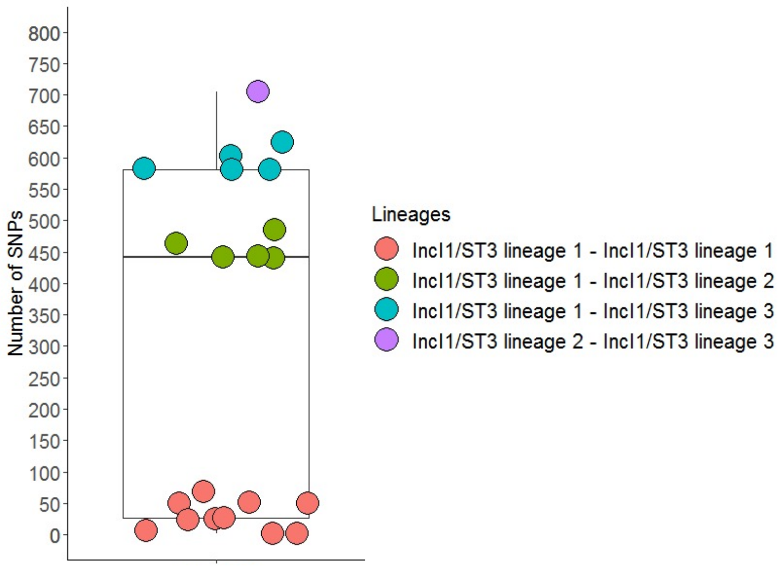


C

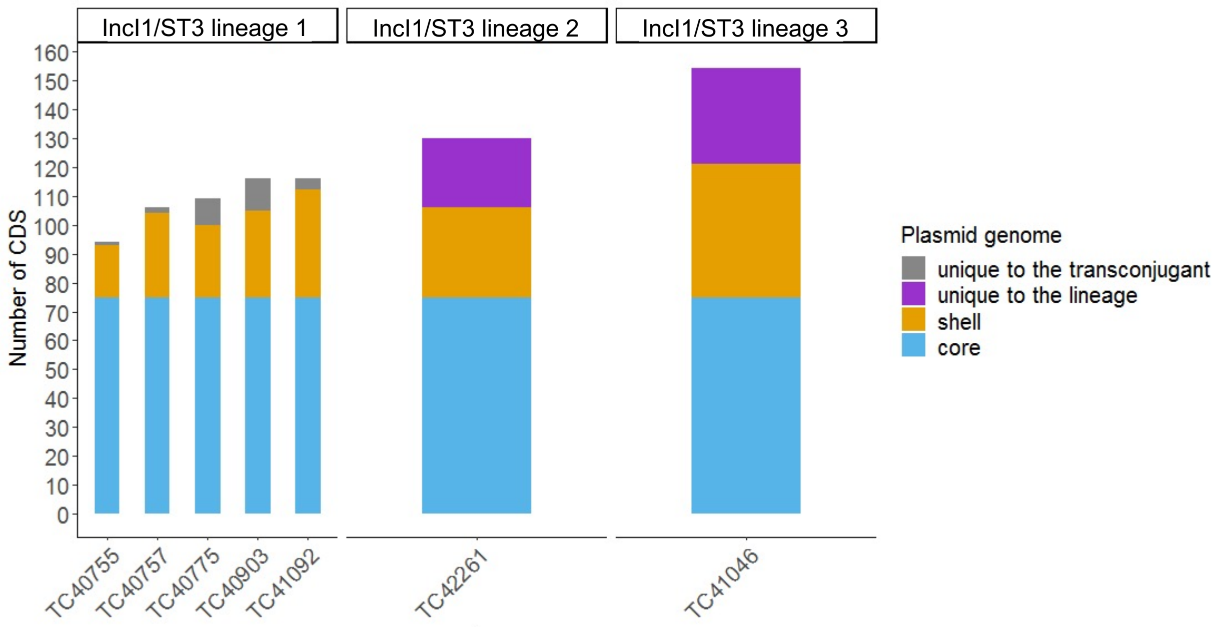

**Fig. S3B. PFGE profiles of the ESBL-producing *E. coli* isolates in farm B.** Pulsed-field gel electrophoresis (PFGE) was performed on all putative ESBL-producing *E. coli* isolates using the restriction enzyme *Xba*I. Comparison of PFGE profiles was done among isolates to discriminate between ESBL-producing *E. coli* strains circulating in the farm and to select one representative of each PFGE profile for genome sequencing. Two isolates were considered to be related to different genotypes if their PFGE profiles differed from at least one band. Isolates having a different status regarding the presence of the *mcr-1* gene compared to their group were additionally selected for genome sequencing. The isolate ID and the sample code (the animal ID followed by the sampling rank) are displayed on the left of the PFGE patterns. On the right, genotypic markers characterized on all isolates, *i.e.* phylogroup, $bla_{CTX\text{-}M}$ enzyme and presence of the *mcr-1* gene are represented. The clone ID and the $bla_{CTX\text{-}M}$ gene molecular support are indicated in front of the isolates that were sequenced. Plasmid sequences were obtained from isolates with clone ID in bold after conjugation with a K-12 strain. Isolates of different clones are colored according to the $bla_{CTX\text{-}M}$ gene molecular support. COL: Colistin, ESC: Extended-Spectrum Cephalosporins.

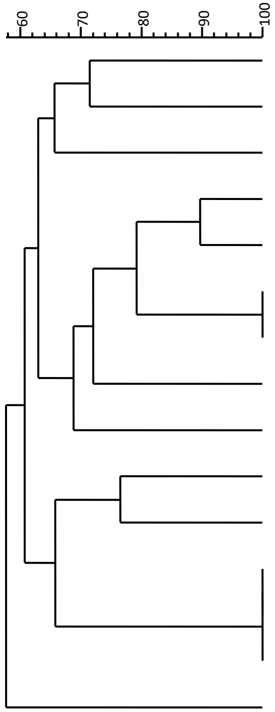


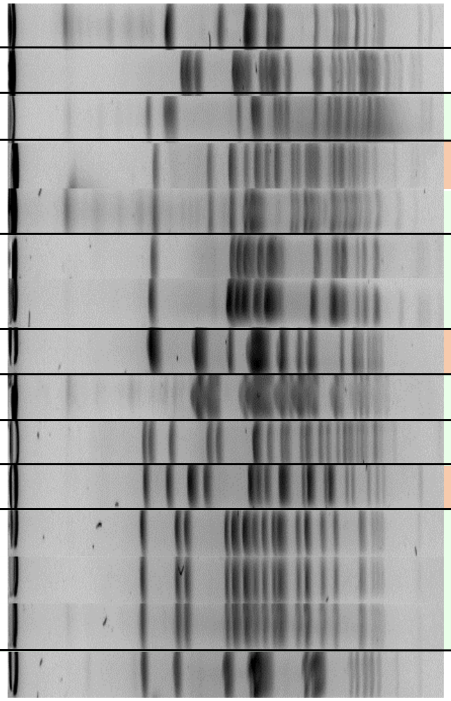

| Isolate ID | Sample code | Phylogroup | Clone ID | ESC-resistance enzyme | COL-resistance enzyme | $bla_{CTX\text{-}M}$ gene location |
|---|---|---|---|---|---|---|
| 40772 | 7695 E0 | A | A ST10/2 O101:H9 *fimH54* clone2 | CTX-M-15 | - | chromosomic |
| 41031 | 7640 E2 | A | A ST1139/958 O8:H9 *fimH54* | CTX-M-14 | - | IncF/F2:A-:B- |
| 40775 | 8660 E0 | A | **A ST10/466 O148:H32 *fimH23*** | CTX-M-1 | - | IncI1/ST3 lineage 1 |
| 40752 | 1005 E0 | A | **A ST301/917 O80:H2 *fimH54*** | CTX-M-1 | - | IncI1/ST312 |
| 40769 | 7640 E0 | A | A ST301/917 O80:H2 *fimH54* | CTX-M-1 | - | IncI1/ST3 (lineage unknown) |
| 40755 | 1601 E0 | A | **A ST167/2 O101:H9 *fimH215*** | CTX-M-1 | - | IncI1/ST3 lineage 1 |
| 40767 | 6205 E0 | A | | CTX-M-1 | - | |
| 40892 | 2222 E1 | B1 | **B1 ST58/24 O8:H25 *fimH34*** | CTX-M-1 | - | IncI1/ST312 |
| 41092 | 1005 E3 | B1 | **B1 ST58/24 O9:H25 *fimH32*** | CTX-M-1 | - | IncI1/ST3 lineage 1 |
| 40757 | 1603 E0 | E | **E ST753/920 O130:H9 *fimH124*** | CTX-M-1 | - | IncI1/ST3 lineage 1 |
| 40759 | 2222 E0 | G | **G ST117/48 O33:H4 *fimH41*** | CTX-M-1 | - | IncI1/ST312 |
| 40765 | 3253 E0 | B1 | | CTX-M-1 | - | |
| 40903 | 3253 E1 | B1 | **B1 ST2522/596 O29:H34 *fimH38*** | CTX-M-1 | - | IncI1/ST3 lineage 1 |
| 41028 | 3253 E2 | B1 | | CTX-M-1 | - | |
| 40762 | 2522 E0 | B1 | **B1 ST162/25 O8:H19 *fimH32*** | CTX-M-14 | - | IncF/F2:A-:B- |
