## Supplementary Figures S5 S6 for "Interplay between bacterial clone and plasmid in the spread of antibiotic resistance genes in the gut: lessons from a temporal study in veal calves"

**Fig. S5A. Distribution of extraintestinal pathogenic virulence genes detected in ESBL-producing *E. coli* clones.** Heatmap of the virulence genes associated with extraintestinal pathogenic *E. coli*, using the VFDB and VirulenceFinder databases. Virulence genes are indicated on the x-axis and are grouped per function encoded. ESBL-producing *E. coli* clones are indicated on the y-axis and are grouped according to the farm in which they were isolated. The clone was defined by a combination of the phylogroup, ST Achtman/ST Pasteur Institute, serotype, and *fimH* allele gene. The number of extraintestinal pathogenic virulence genes detected in each clone is displayed on the right.

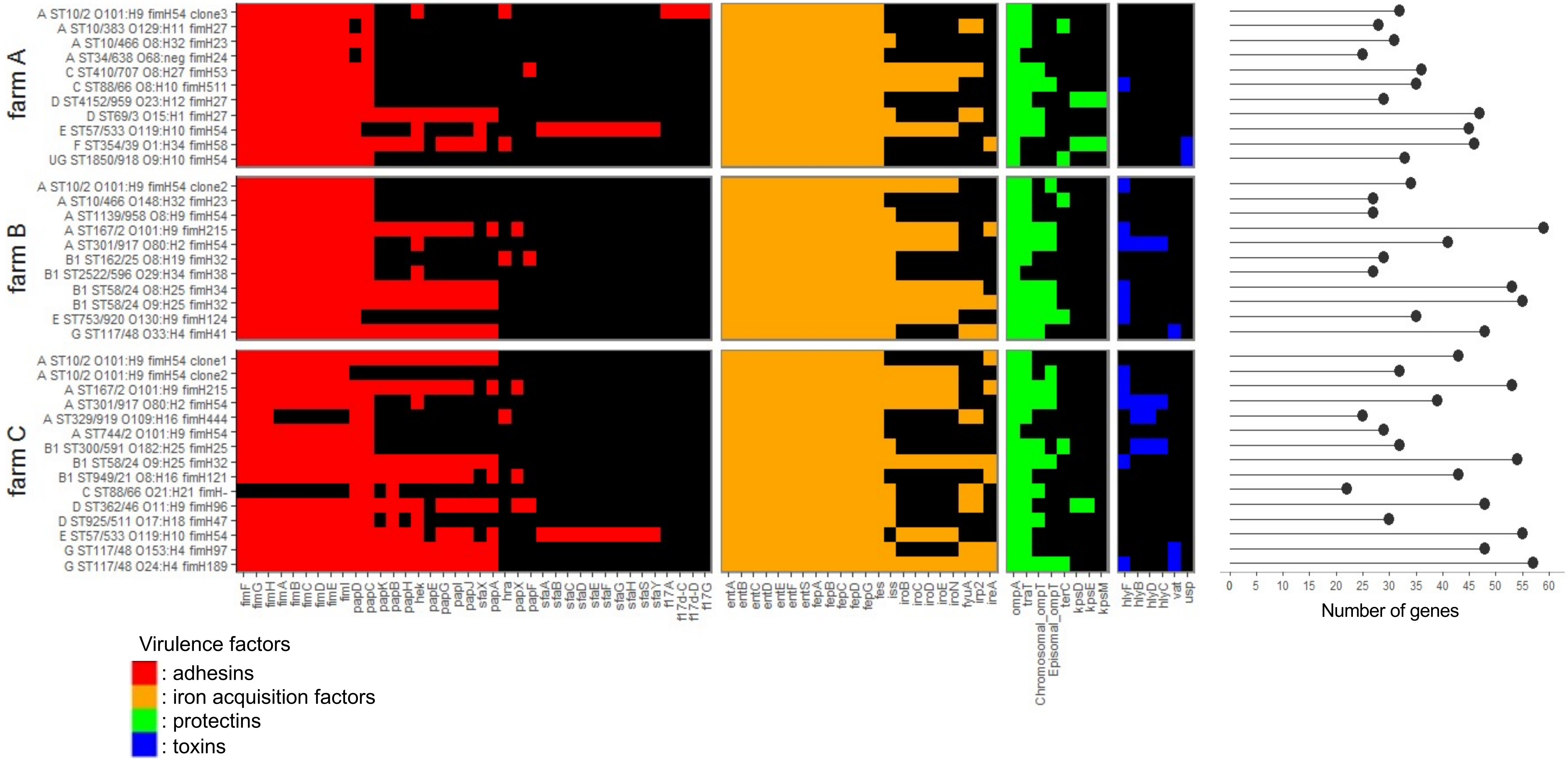

**Fig. S5B. Distribution of intestinal pathogenic virulence genes detected in ESBL-producing *E. coli* clones.** Heatmap of the virulence genes associated with intestinal pathogenic *E. coli*, using the VFDB and VirulenceFinder databases. Virulence genes are indicated on the x-axis and are grouped per function encoded. ESBL-producing *E. coli* clones are indicated on the y-axis and are grouped according to the farm in which they were isolated. The clone ID is a combination of the phylogroup, ST Achtman/ST Pasteur Institute, serotype, and *fimH* allele gene. The number of intestinal pathogenic virulence genes detected in each clone is displayed on the right.

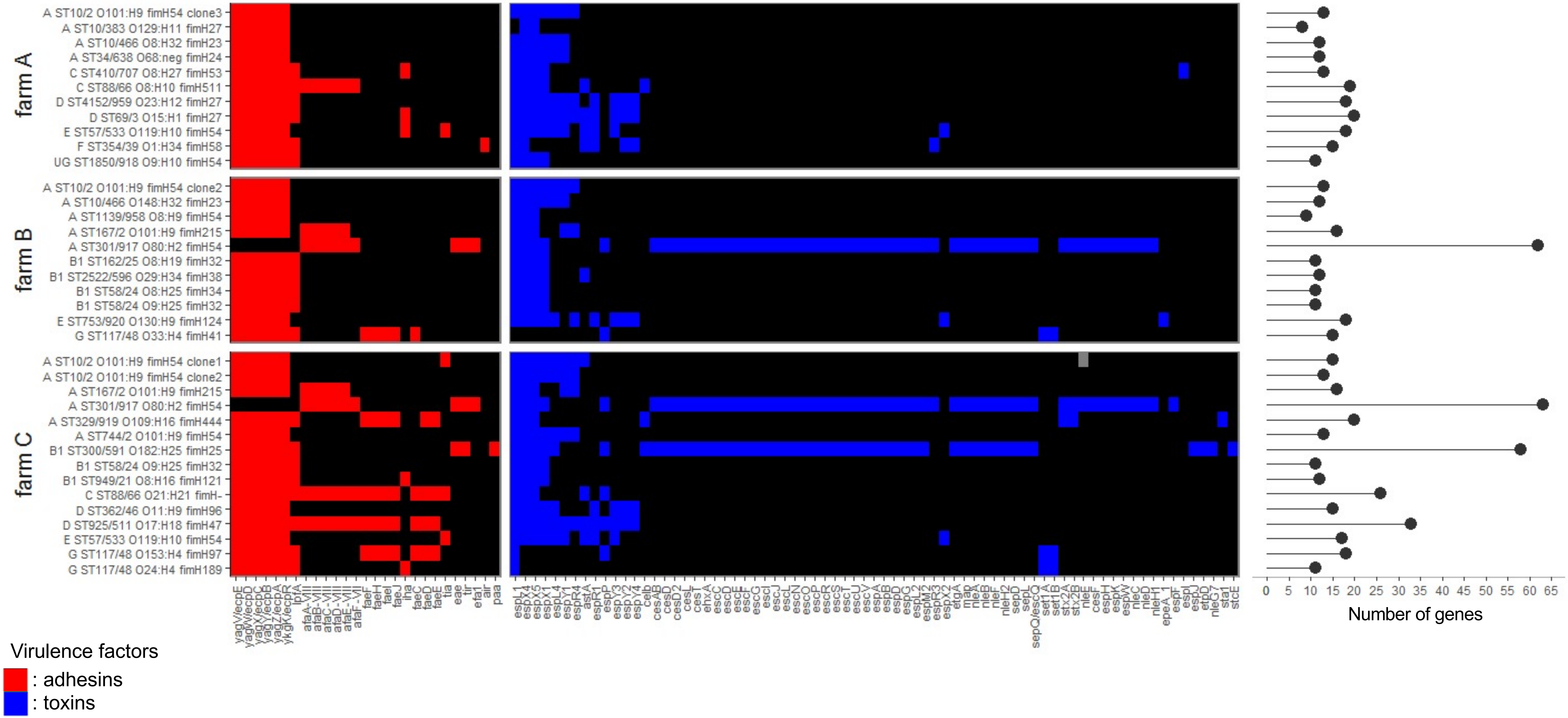

**Fig. S5C. Distribution of pathogenic virulence genes associated with both extraintestinal and intestinal pathogenicity, and bacteriocins detected in ESBL-producing *E. coli* clones.** **H**eatmap of the virulence genes associated with both extraintestinal and intestinal pathogenic *E. coli*, using the VFDB and VirulenceFinder databases, and of the bacteriocins detected in the isolates sequenced. Virulence genes are indicated on the x-axis and are grouped per function encoded. ESBL-producing *E. coli* clones are indicated on the y-axis and are grouped according to the farm in which they were isolated. The clone ID is a combination of the phylogroup, ST Achtman/ST Pasteur Institute, serotype, and *fimH* allele gene. The number of virulence genes detected in each clone is displayed on the right.

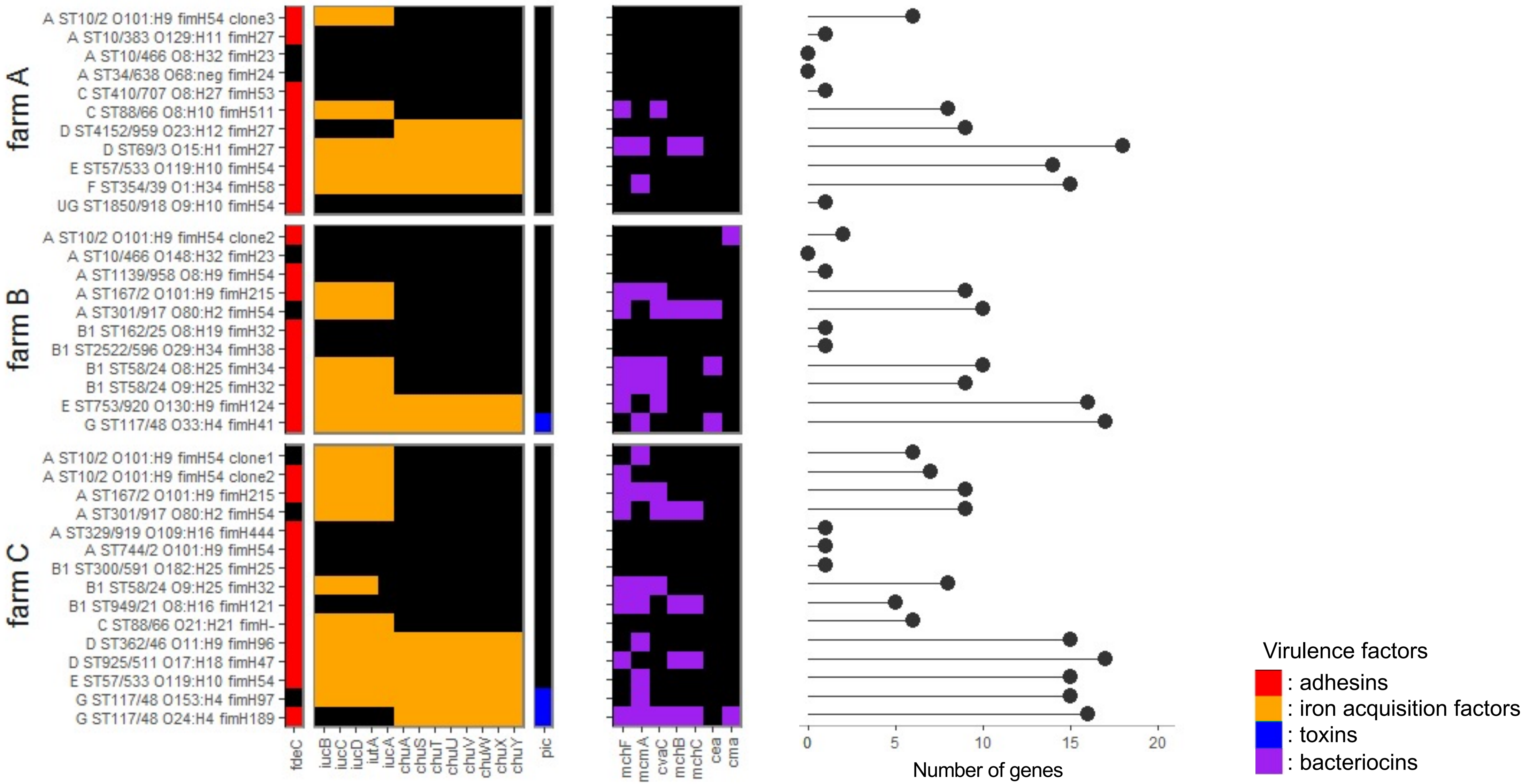

**Fig. S6A. Linear map of the chromosomal environment of *$bla_{CTX\text{-}M\text{-}15}$* gene and *mcr-1* gene.** The chromosomal environment of the *$bla_{CTX\text{-}M\text{-}15}$* gene found in the clone 'A ST10/2 O101:H9 *fimH54* clone2' isolated in farms B and C (isolate IDs 40772 and 41054, respectively) is represented on the top of the figure. The chromosomal environments of the *mcr-1* gene in clones 'F ST354/39 O1:H34 fimH58' in farm A (isolate ID 40808) and 'G ST117/48 O24:H4 fimH189' in farm C (isolate ID 41301), are represented on the middle and the bottom of the figure, respectively. The antibiotic resistance genes are indicated by red arrows. Open reading frames are shown as arrows indicating the direction of transcription (dark blue, plasmid transfer; red, resistance; pink, mobile elements). The name of genes is indicated within the arrows. The genome references against which contigs containing the genes were blasted are indicated below the maps.

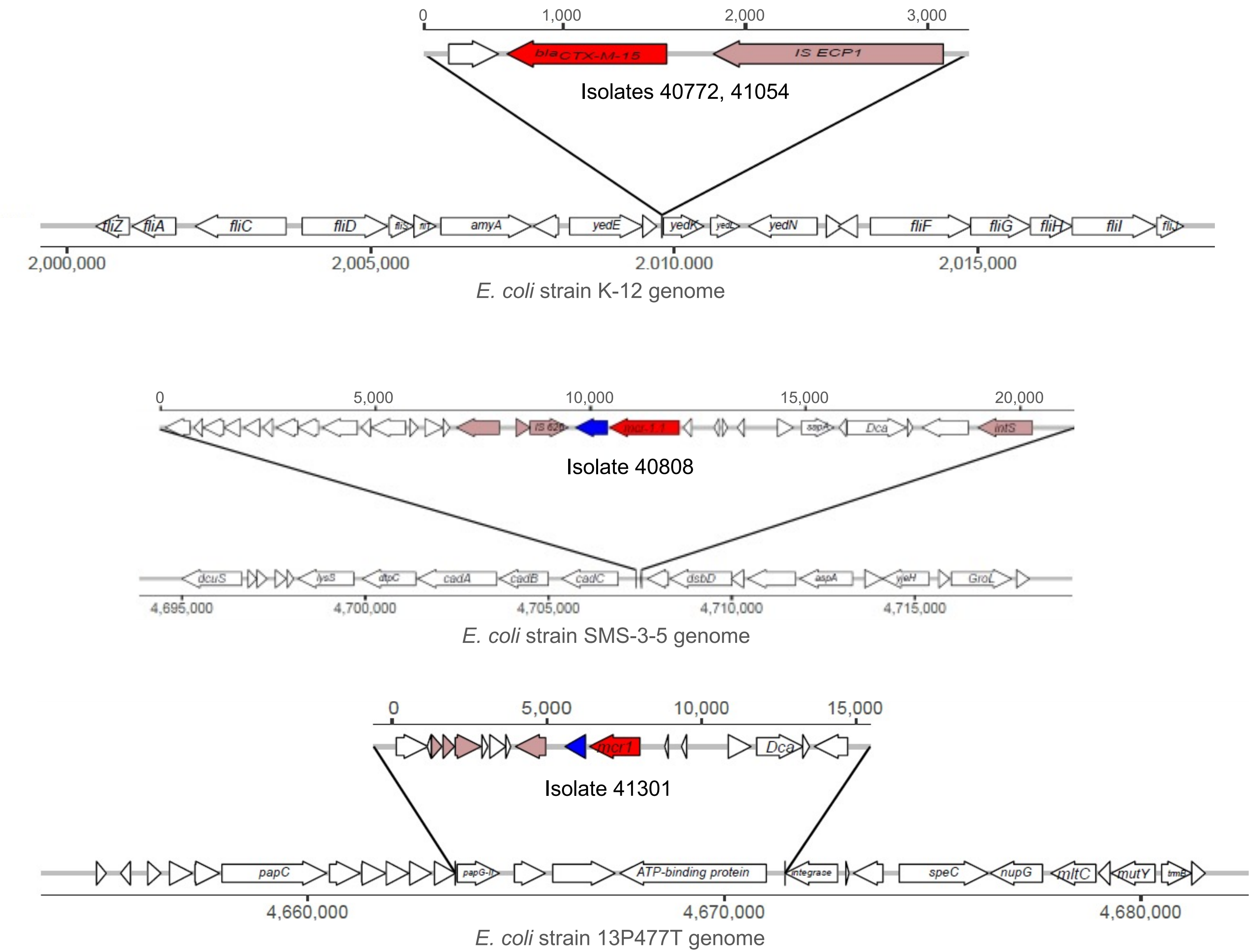

**Fig. S6B. Circular maps of prevalent plasmids carrying $bla_{CTX-M}$ genes and *mcr-1* gene.** The genome of transconjugants (TC) carrying plasmid-borne ESBL-encoding genes or *mcr-1* gene were sequenced to characterize the molecular environment of these antibiotic resistance genes. Plasmids were identified using the PlasmidFinder database and reference genomes were found by blasting contigs against MaGe or Refseq databases. The map and size of plasmids IncI1/ST312, IncF/F2:A-:B-, and IncX4 are represented on the left, the middle and the right of the plot, respectively. Open reading frames are shown as boxes colored according to the function of their product. ESBL-encoding genes ($bla_{CTX-M-1}$ and $bla_{CTX-M-14}$ genes) and *mcr-1* gene are indicated in red on the map. Single Nucleotide Polymorphisms (SNPs) detected in the plasmidic contigs of the TC are represented by black lines. The donor strain ID of each TC is given below the figure, and the farm in which it was isolated is provided between brackets.

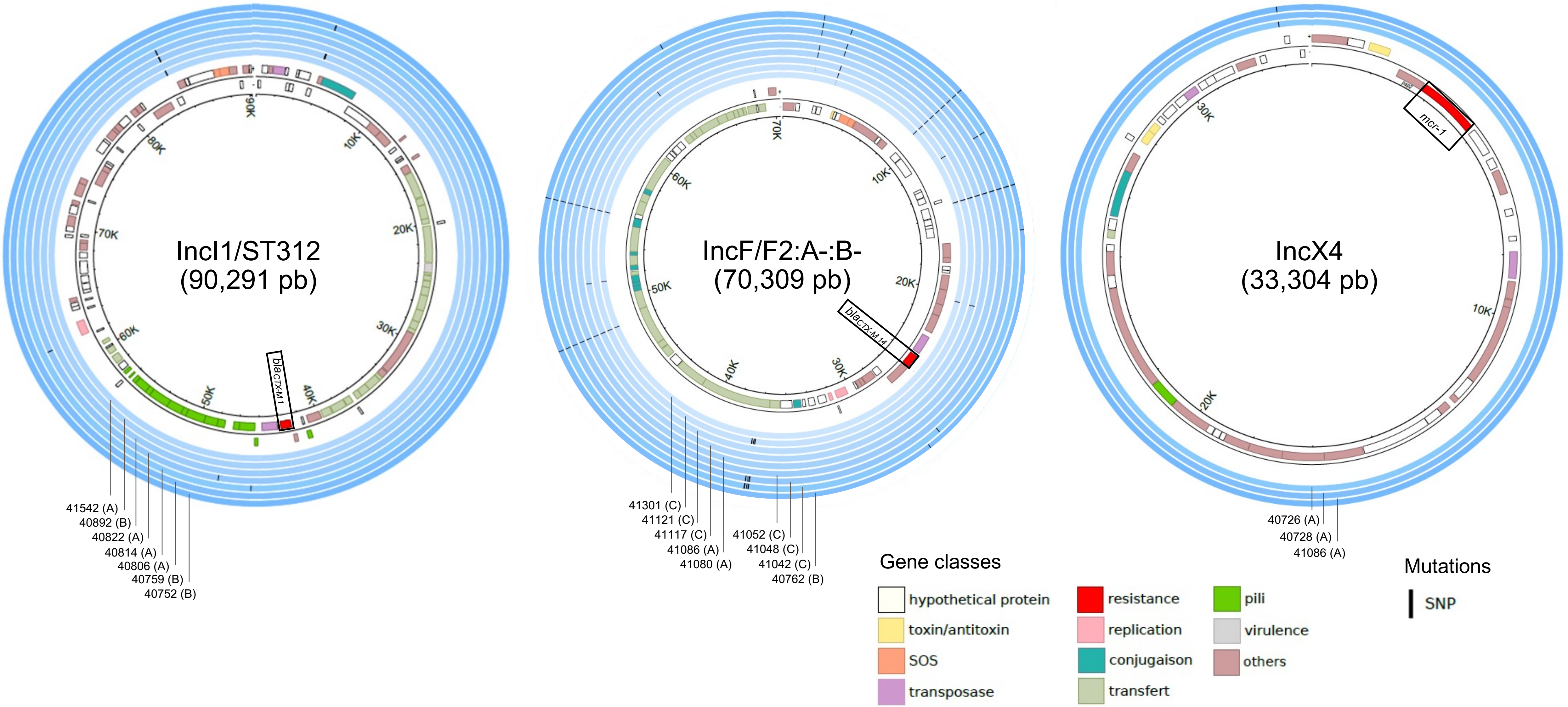

**Fig. S6C. Circular maps of plasmids carrying $bla_{CTX-M}$ genes.** The genome of transconjugants (TC) carrying plasmid-borne ESBL-encoding genes or *mcr-1* gene were sequenced to characterize the molecular environment of these antibiotic resistance genes. Plasmids were identified using the PlasmidFinder database and reference genomes were found by blasting contigs against MaGe or Refseq databases. The map and size of plasmids IncI1/ST3, IncF/F59:A-:B-, and IncK are represented on the left, the middle and the right of the plot, respectively. Open reading frames are shown as boxes colored according to the function of their product. ESBL-encoding genes ($bla_{CTX-M-1}$ and $bla_{CTX-M-14}$ genes) and *mcr-1* gene are indicated in red on the map. Single Nucleotide Polymorphisms (SNPs) and deletions detected in the plasmidic contigs of the TC are represented by black and white lines, respectively. The donor strain ID of each TC is given below the figure, and the farm in which it was isolated is provided between brackets. For IncI1/ST3 plasmid, the lineage of each plasmid sequenced is indicated below the figure.

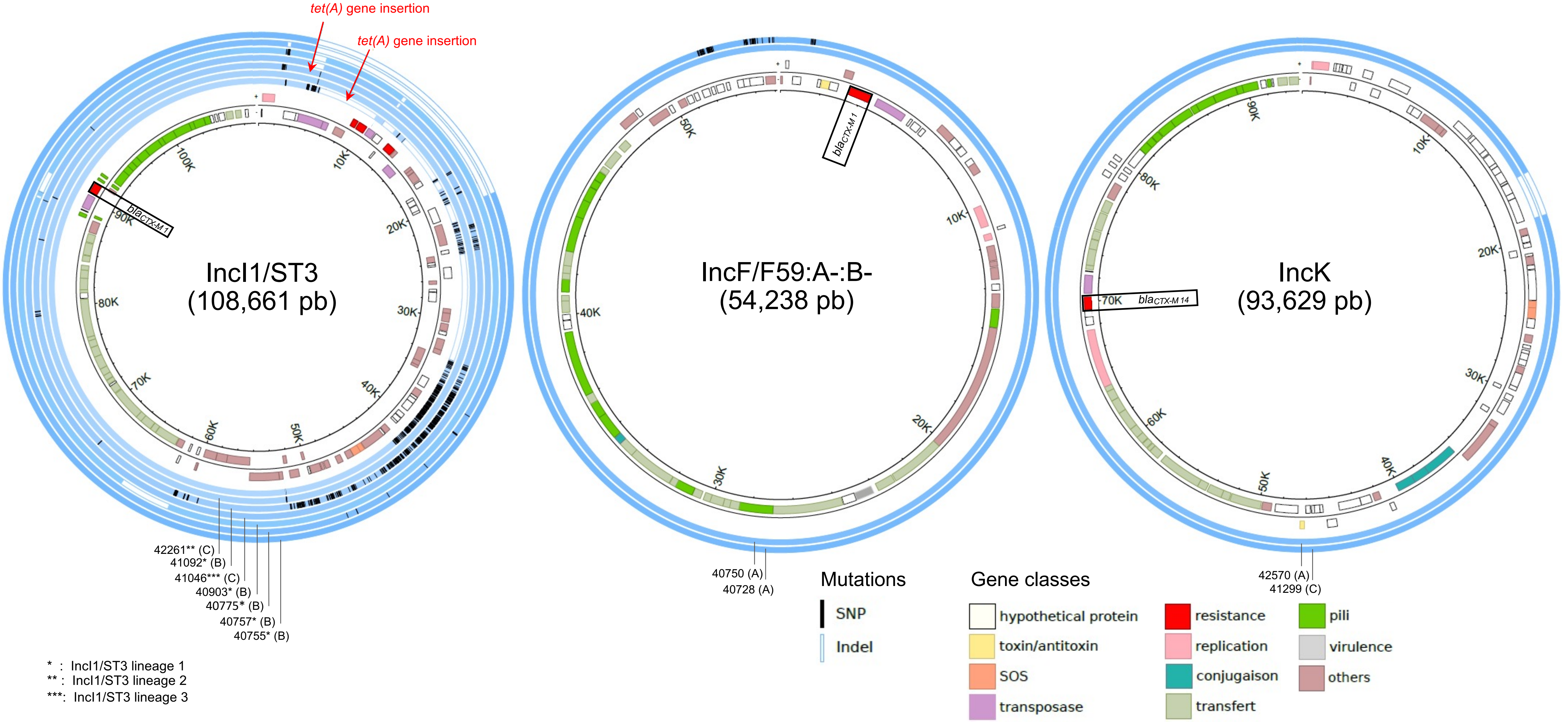


* : IncI1/ST3 lineage 1
** : IncI1/ST3 lineage 2
***: IncI1/ST3 lineage 3
