## Supplementary Text for "Interplay between bacterial clone and plasmid in the spread of antibiotic resistance genes in the gut: lessons from a temporal study in veal calves"

**Supplementary Material and methods**

**Study design and animal selection**

Sampling began upon the arrival of batches of new calves in October and November 2015. Each farm is counting 250-300 animals per batch, a batch being defined as a group of calves entering the farm at the same time and reared together until slaughter.

**Sampling and antibiotic data collection**

Antibiotics were always used at therapeutic doses and administered orally in water or milk replacer. All treatments upon entry to the fattening farms were set up to prevent gastrointestinal disorders, whereas treatments during the course of fattening were used to treat respiratory diseases.

All calves received antibiotics more than once and calves from farms A and C received several consecutive antibiotic treatments during the first month (Fig. 1). On farm A, calves received a 10-day course of colistin and sulfonamides (first day to day 10), followed by another 10-day course of tetracycline (day 11 to day 20). Later during fattening, they received a one-day treatment of doxycycline on day 53 and a one-day treatment of amoxicillin on day 135. On farm B, calves received a one-day treatment of doxycycline and erythromycin on day 26, a one-day treatment of tetracycline on day 90, and a one-day treatment of amoxicillin on day 101. On farm C, calves received a six-day course of sulfonamides and trimethoprim (from day 3 to day 8) and a seven-day course of tetracycline (day 10 to day 16). They also received a five-day course of doxycycline (day 20 to day 24) and four-day courses of spiramycin (day 25 to day 28) and tetracycline (day 80 to day 83).

**Quantification of *bla*_CTX-M_ copy number in the fecal samples**

The sequences of forward (5’-GGA ATC TGA CGC TGG GTA AA-3’) primer, reverse (5’-GGT TGA GGC TGG GTG AAG TA-3’) primer, and probe (5’-HEX-ACT ATG GCA CCA CCA ACG AT-BHQ-1-3’) for detecting *bla*_CTX-M_ group 1 were obtained from (1). The sequences of the forward (5’-GGT GAT GAA CGC TTT CCA AT-3’) and reverse (5’-TCA ATT TGT TCA TGG CGG TA-3’) primers and probe (5’-6-FAM-CAG AGT GAA ACG CAA AAG CA-BHQ-1-3’) for detecting *bla*_CTX-M_ group 9 were also obtained from (1). Each DNA sample (approximately 20 ng) was added to 25 μl PCR mixture containing 12.5 μl TaqMan Universal PCR master mix II 2X (Applied Biosystems, Life Technologies, Carlsbad, California, USA), 200 nM of each primer, and 50 nM of each fluorescent probe, as previously described in (1).

Ten-fold serial dilutions of extracted plasmid DNA containing either *bla*_CTX-M-15_ or *bla*_CTX-M-9_ genes from a recipient *E. coli* strain K-12 and using QIAprep spin miniprep kit (QIAGEN, Venlo, Netherlands) were used to build standard curves for *bla*_CTX-M_ group 1 and *bla*_CTX-M_ group 9, respectively (*bla*_CTX-M-15_ carrying plasmid ID ‘RCS102’, GenBank accession no ‘LT985213’; *bla*_CTX-M-9_ carrying plasmid ID ‘RCS51’, GenBank accession no ‘LT985252’, (2)).

**Assessment of the prevalence of *E. coli* among *Enterobacteriaceae* in calves’ feces**

We considered that the colonies were mainly from *E. coli* species, as it is quantitatively a major member of the *Enterobacteriaceae* family in cattle’s feces. We verified this assumption by plating three randomly selected swabs from distinct calves of our study on Drigalski plates. We determined the species of 50 colonies per plate using MALDI-TOF and Clermont typing PCR (3). All colonies were confirmed to be *E. coli*.

**Selection of ESBL-producing *E. coli* isolates**

Upon their arrival at the Anses lab, swabs were plated on selective ChromID ESBL agar for the detection of ESBL-producing *E. coli,* as well as on MacConkey agar as a control of growth (data not shown).

**Antimicrobial susceptibility testing of ESBL-producing *E. coli* isolates**

A total of ten non-β-lactam - colistin, tetracycline, kanamycin, gentamicin, streptomycin, florfenicol, sulfonamides, trimethoprim, nalidixic acid, and enrofloxacin - antibiotics of veterinary and human interests was tested. The interpretation was based on the breakpoints provided by the Antibiogram Committee of the French Society of Microbiology for *E. coli* (<http://www.sfm-microbiologie.org>). Susceptibility to 16 β-lactams (amoxicillin, piperacillin, ticarcillin, amoxicillin-clavulanic acid, piperacillin-tazobactam, ticarcillin-clavulanic acid, cefalotin, cefuroxime, cefotaxime, ceftiofur, ceftazidime, cefoxitin, cefepime, cefquinome, aztreonam, and ertapenem) was also tested, and ESBL production was confirmed by the double-disc synergy test. *E. coli* ATCC 25922 strain was used as the quality control. The MIC to colistin was determined using the broth microdilution method.

**Genotypic discrimination of ESBL-producing *E. coli* isolates**

Two isolates were considered to be different strains if their PFGE profiles differed from at least one band. The delineation of strains by PFGE typing was confirmed by multilocus variable number tandem repeat analysis (MLVA) (4). MLVA typing also allowed typing isolates for which PFGE yielded a smearing profile.

**Whole genome sequencing (WGS) of ESBL-producing *E. coli* isolates**

*Genome sequencing of the* E. coli *isolates*

DNA was simultaneously fragmented and tagged with sequencing adapters in a single step using Nextera DNA library prep kit (Illumina, San Diego, California, USA). Multiplexing barcode sequences were added to the libraries by limited-cycle PCR using Nextera technology. Barcoded libraries were sequenced on an Illumina HiSeq 4000 according to the manufacturer’ specifications to generate paired-end reads of 100 base pairs (bp) in length.

*First step bioinformatics process (quality check and assembly)*

The quality of the reads was checked using FASTQC (version 0.11.6) (5). The reads were assembled to make contigs using SPAdes (6). The depth of sequencing was assessed for all the genomes sequenced by multiplying the number of reads by their length and dividing it by the size of the *E. coli* strain ED1a genome, which is 5,209,548 base pairs (7).

*Phylogenetic analyses and clone definitions*

For isolates from different haplogroups, whether or not from the same phylogroup, the mean reached 15,366 (± 8,572) SNPs and 48,147 (± 13,911) SNPs, respectively ( Fig. S1A). The mean number of SNPs in the core genome between two isolates of the same haplogroup dropped to 457 (± 748). The number of SNPs in the core genome between two isolates from the same haplogroup was significantly inferior to the number of SNPs between isolates of different haplogroups (Wilcoxon tests, adjusted p <10^-10^), which validate our choice regarding the combination of genotypic markers we used to group isolates.

A threshold of 100 SNPs was chosen according to a gap ranging from 100 to 1,000 SNPs to delineate clones from the same haplogroup (Fig. S1). All the haplogroups that had more than one sequenced isolate had less than 100 SNPs of difference between them, except for the haplogroup ‘ST10/2 O101:H9 *fimH54*’. In this haplogroup, four isolates had been sequenced and only two of them had a number of SNPs inferior to 1,000 SNPs (Fig. S1B). Thus, this pair of isolates was considered to be part of a distinct clone from the other isolates, even though they had a different PFGE profile (Fig. S1C). This haplogroup was composed of the three clones ‘ST10/2 O101:H9 clone1’, ‘ST10/2 O101:H9 clone2’, ‘ST10/2 O101:H9 clone3’. Isolates of the same clone differed in mean by 18 SNPs (± 20) on their core genome, the maximum being 54 SNPs for the clone ‘ST301/917 O80:H2’ (Fig. S1D).

A maximum likelihood phylogenetic tree was built using FastTree 2 from Harvest (8, 9). The tree was rooted on the *E. coli* strain ED1a, from the B2 phylogroup. The tree structure was visualized with iTOL (version 3) (10). Isolates were identified by their ID followed by their clone’s name.

*Resistome and virulome analyses*

Contigs were screened for antibiotic resistance and virulence genes using the software Abricate 0.8.1 (using parameters minID 80%, minCOV 90%) (11). The database ResFinder (version 3.0) was used to detect antibiotic resistance genes (12). A custom database mixing VFDB (13), Virulence Finder (version 2.0) (14), and specific genes from extraintestinal pathogenic *E. coli* were used to detect the virulence genes. The localization of the genes on chromosome or plasmid was determined using the software Centrifuge (version 1.0.3) (15) within the PlaScope workflow (16). Quinolone resistance point mutations were also searched using PointFinder with standard parameters (September 2018) (17, 18). The full list of antibiotic resistance genes and virulence genes of the strains sequenced are presented in Table S1A, along with their genome accession numbers.

In summary, the available genomic data for each isolate sequenced are the phylogenetic group, the STs according to the Warwick scheme and Pasteur institute scheme, the serotype and the *fimH* gene allele, the number of SNPs in the core genome compared to the other isolates (all these markers being used in combination to delineate clones), the resistome and the virulome contents.

**Conjugation and plasmid sequencing**

*Selection of the plasmids to be sequenced*

All transconjugants (TC) were controlled by PCR using adequate primers for the detection of either the *bla*_CTX-M_ gene (19) or the *mcr-1* gene (20). Only TC presenting a unique plasmid, as assessed by S1-PFGE, were further characterized.

*Phylogenetic analyses and plasmid lineage definition*

The delineation of plasmids in clones was explored for plasmids that had more than one isolate sequenced, *i.e.* IncF/F2:A-:B- plasmids (nine reconstructed genomes), IncI1/ST3 plasmids and IncI1/ST312 plasmids (both seven reconstructed genomes), IncX4 plasmids (three reconstructed genomes), IncF/F59:A-:B- plasmids, IncK plasmids (both two reconstructed genomes). The mean number of SNPs in the core genome between two plasmids of the same group was 83 (± 187). A threshold of 100 SNPs was chosen according to a gap ranging roughly from 100 to 400 SNPs (Fig. S2A). All the sequenced plasmids had less than 100 SNPs of difference between their isolates, except for the IncI1/ST3 plasmid (Fig. S2A).

In IncI1/ST3 plasmid, seven isolates had been sequenced and two of them had a number of SNPs superior to 100 SNPs with all the other isolates, as well as between them (Fig. S2B). Thus, these isolates were considered to be part of distinct lineages from the other isolates. This plasmid was composed of three lineages named ‘IncI1/ST3 lineage1’, ‘IncI1/ST3 lineage2’, and ‘IncI1/ST3 lineage3’. This delineation was in accordance with the variable gene pool of the different lineages (Fig. S2C). Even if more than 50% of the gene content was shared among different lineages, lineage 2 and lineage 3 had 24 and 33 genes not present in the other lineages, respectively.

*Identification of* bla*_CTX-M_ gene carrying-plasmids and* mcr-1 *gene carrying-plasmids in the collection of isolates*

For 14 clones (out of 15) that were isolated one time, we sequenced the plasmid carrying the *bla*_CTX-M_ gene. For 14 clones (out of 16) that had a plasmid-borne *bla*_CTX-M_ gene and were isolated several times, we sequenced the plasmid carrying the *bla*_CTX-M_ gene in one (or two) isolate(s). The plasmid identified in a clone as carrying the *bla*_CTX-M_ gene was detected in 100% of the isolates of 11 clones (representing a total of 92 isolates), and the majority of the isolates for one of them (21/22 isolates). The plasmid identified in a clone as carrying the *bla*_CTX-M_ gene was not detected in a few isolates: 2 isolates for the ‘ST301/917 O80:H2’ clone (67% of the isolates of this clone) and 11 isolates for the ‘ST744/2 O101:H9’ clone (50% of the isolates).

For four clones (out of seven) in which a plasmid-borne *mcr-1* gene was detected, we sequenced the plasmid carrying it. The plasmid identified in a clone as carrying the *mcr-1* gene was detected in 100% of the isolates of this clone in which the *mcr-1* gene had been previously detected by PCR.

**Statistical analyses**

We used linear regression analysis with 1,000 permutations to test the effect of the level of excretion of ESBL-producing *E. coli* at day 7 on the number of positive samples during the fattening. Data were initially explored at group level using Kruskal-Wallis tests to assess difference in the number of ESBL positive samples per calf between farms and between the groups of calves as classified at day 7 (ESBL-producing *E. coli* high-level carrier”, “low-level carrier” or “ESBL-producing *E. coli*-free”). These variables were subjected to linear regression analysis with permutations if significant differences were observed between groups in the initial tests. After visual inspection of the data, an interaction term between the two variables was also tested. Tests were performed using 1,000 permutations. The likelihood ratio test (LRT) was used to compare the fit between the candidate models with and without the interaction term.

The Spearman correlation test was used to look for an association between the number of calves colonized by a clone and its period of detection in a farm. For the five clones that were detected in several farms, we considered the diffusion and the persistence in the farm in which both of these parameters were the highest. The Spearman correlation test was also used to look for an association between these two variables and the antibiotic coresistance score of each clone. We chose among its isolates the highest maximum score of a clone to search for this putative association.

The Roary software (21) was used to calculate the pan-genome of the set of clones and the Scoary software (22) was used to score genes for associations with persistence and diffusion traits (standard parameters were used). As multiple tests were performed, Bonferroni correction was used to adjust p-values.

Reconstructed plasmids were visualized using the R package circlize (version 0.4.7) (23). Chromosomal insertions of the *mcr-1* gene and the *bla*_CTX-M_ genes were visualised with the packages ggplot2 and gggenes (version 0.4.0) (24).

**Supplementary Results**

**Quantification of the excretion of ESBL-producing *E. coli* and of *bla*_CTX-M_ genes**

Three ESBL-producing *E. coli* carriage patterns were found: calves in which ESBL-producing *E. coli* were never detected, calves that carried ESBL-producing *E. coli* during the first half of fattening and then were cleared, and calves which carriage persisted over the fattening (Fig. 2A). These patterns were not equally distributed among farms. In farms A and B, calves were mainly transitory carriers, even though the excretion was shorter in farm B compared to farm A. In farm A, all calves had a long-lasting excretion of ESBL-producing *E. coli* during the first two months (Fig. 2A), followed by a general clearance phenomenon during the third month. In farm B, no ESBL-producing *E. coli* was detected from day 63 until the end of fattening (Fig. 2A). Calves that never carried ESBL-producing *E. coli* were only found in farm B. In farm C, all calves were persistent carriers, although the carriage was unsteady (Fig. 2A).

The number of positive samples per calf was significantly associated with the farm (Kruskal-Wallis test, p= 3 x 10^-7^) and the level of excretion at day 7 (Kruskal-Wallis test, p= 2 x 10^-3^) and so the two variables were used as covariables in the linear regression. The interaction term didn’t provide a significant additional explanatory power (LRT between the two candidate models, p= 0.8) and thus the model with no interaction term was preferred.

The *bla*_CTX-M_ genes were detected in only one sample from which no colony had grown on ChromID ESBL agar (0.5% of the 186 negative samples tested). As the two detection methods were done on duplicate swabs, it highlights the reliability of rectal swabbing to monitor the carriage of ESBL-producing *E. coli* in calves. The level of excretion of *bla*_CTX-M_ genes in feces was high during the first two months in farm A, reaching a mean of 7.0 log_10_(copies */* g), 4.9 log_10_(copies */* g) and 5.4 log_10_(copies */* g) at day 7, day 21 and day 49, respectively (Fig. 2B). In farm B, the *bla*_CTX-M_ genes were detected only in 6 calves (out of ten) at day 7 and in no calves at day 21 (Fig. 2B). The maximum of excretion was on day 7, with a mean of 2.7 log_10_(copies */* g). In farm C, calves experienced two peaks of excretion, at days 35 and 119, with a mean of 5.3 log_10_(copies */* g) and 2.8 log_10_(copies */* g), respectively (Fig. 2B).

**Characterization of ESBL-producing *E. coli* isolates**

The *bla*_CTX-M-1_ gene was the most prevalent *bla*_ESBL_ gene among isolates in farm A (64/84 isolates, 76.2%) and in farm B (12/15 isolates, 80.0%). The *bla*_CTX-M-1_ and *bla*_CTX-M-14_ genes were found in equal proportions in farm C (36/74 for both, 48.6%).

Most of the ESBL-producing *E. coli* isolates were resistant to tetracycline and sulfonamides (Table S3A). Some co-resistances were found in more than 80% of the isolates of a given farm, such as kanamycin in farm B, streptomycin in farms B and C (more than 85% in both farms), and trimethoprim in farms A and B (more than 80% in both farms, Table S3A). Resistances to gentamicin, florfenicol and quinolones were found in half or less of the isolates in each farm (Table S3A). Seven ESBL-producing isolates (4.6%) in farm A and one in farm C were resistant to all the antimicrobials tested.

**Molecular characterization of the ESBL-producing *E. coli* clones**

*Phylogenetic characterization of the ESBL-producing E. coli isolates*

The mean depth of sequencing was 29.0 (±13.6, Table S1A). The depths ranged from 19.4 to 113.6. A phylogenetic tree was reconstructed from 200,875 SNPs of the 3,003 genes that composed the core genome of this set of sequenced genomes. The pan-genome was composed of 15,145 genes.

The *bla*_ESBL_ genes found were *bla*_CTX-M-1_ (112/173 isolates, 64.7%), *bla*_CTX-M -14_ (58/173, 33.5%), and *bla*_CTX-M-15_ (3/173, 1.7%). The *bla*_CTX-M-1_ gene and *bla*_CTX-M-14_ gene were detected in all farms. The *bla*_CTX-M-15_ gene was found in farm B and in farm C. Most ESBL-producing isolates in farm A carried the *mcr-1* gene (52/84 isolates, 61.9%), whereas only one isolate carrying it was found in farm C, representing a total of 31.2% of the ESBL-producing isolates. The *mcr-1* gene was not detected in any isolate from farm B.

*Antibiotic resistance gene content*

The gene *tet(A)* was found in most of the clones (8/11 clones in farm A, 11/11 clones in farm B and 13/14 clones in farm C, Fig. S4C), and hence was mostly responsible for tetracycline resistance observed in all the isolates (Table S3A, Fig. S4C). Regarding resistance to sulfonamides, the genes *sul1* and *sul2* were highly prevalent in clones found in farm A (both 6/11 clones, Fig. S4C), while the *sul3* gene was found in the clone with the highest number of isolates found in this farm (‘ST10/466 O8:H32’, Fig. S4C). In farm B, the *sul2* gene was present in most of the clones (8/11, Fig. S4C). In farm C, the genes *sul1* (10/14 clones) and *sul2* (8/14 clones) were the most prevalent sulfonamides resistance genes and were found in ‘ST 329/919 O109:H16’ and ‘ST744/2 O101:H9’, the two most isolated clones in this farm (Fig. S4C). The genes *aph(6)-Id, aph(3’’)-Ib, aph(3’)-Ia* were the most prevalent aminoglycosides resistance genes in all farms. The high levels of trimethoprim resistance in farm A (Table S3A) were mainly due to the *dfrA1* gene, which was found in five clones (Fig. S4C), and *dfrA12*, which was found in ‘ST10/466 O8:H32’ (Fig. S4C). In farms B and C, the *dfrA5* gene, and *dfrA17* gene were found in several clones with high numbers of isolates (Fig. S4C).

Twelve clones had mutations in quinolone targets, six of them conferring resistance to enrofloxacin. These six clones had one or two mutations in the *gyrA* gene, and one or two mutations in the *parC* gene (2/11 in farm A, 1/11 in farm B, 4/14 in farm C, Table S3B). Of note, the clone ‘ST744/2 O101:H9’, one of the most isolated clones in farm C (23/74 isolates, 31.1%) was found with two mutations in both the *gyrA* gene (positions S83 and D87) and the *parC* gene (positions S80 and A56).

**Molecular characterization of the ESBL-producing *E. coli* clones**

*Phylogenetic characterization of the ESBL-producing E. coli isolates*

The 32 clones, including four clones with two representatives differing by their accessory genome, are presented in Figure S7. The number of clones identified in farms A, B and C was 11, 11 and 14 clones, respectively. The ST10, which is very prevalent in farm animal settings, was found in all farms and was highly prevalent in farm A, as it was found associated with three clones: ‘ST10/466 O8:H32’ (28/84 isolates, 33.3%), ‘ST10/2 O101:H9 clone3’ (6/84, 7.1%) and ‘ST10/383 O129:H11’ (3/84, 3.6%, Fig. 3A, Fig. 3B, Fig. 3C).

**Characterization of the molecular environments of *bla*_CTX-M_ and *mcr-1* genes**

*Plasmid-borne genes*

The *bla*_CTX-M-1_ gene and *bla*_CTX-M-14_ gene were carried on plasmids. Plasmids carrying *bla*_CTX-M-1_ gene found in transconjugants were composed of three distinct lineages of the IncI1/ST3 incompatibility group, the IncI1/ST312 incompatibility group and the IncF/F59:A-:B- incompatibility group. In a given clone, a *bla*_CTX-M_ gene was found on distinct molecular support only for the clone ‘ST301/917 O80:H2’, in which it was localized on a IncI1/ST3 plasmid and the IncI1/ST312 plasmid. Plasmids carrying *bla*_CTX-M-14_ gene found were IncF/F2:A-:B-, IncF/F2:A-:B42, IncF/F2:A-:B49, IncI1/ST80 and IncK plasmids. No other resistance genes were found in these plasmids except for the IncI1/ST3 lineage 1 plasmid, in which *aadA5, dfrA17, sul2* and *tet(A)* were found in some clones (Table S1B).

The *mcr-1* gene was mostly carried on a plasmid, even though a chromosomal insertion was found in two isolates, one from farm A and one from farm C. Plasmids carrying the *mcr-1* gene found in transconjugants were IncX4 and IncHI2 plasmids.

*Chromosomally-encoded genes*

The *bla*_CTX-M-15_ gene was detected on the chromosome of the clone ‘ST10/2 O101:H9 clone2’ in farms B and C. It was found at the same position in the chromosome in each of the isolates (Fig. S6A), flanked by an ISEcp1 insertion sequence and near from the *fliAZY* and *fliGHIJK* loci involved in flagellar synthesis.

The *mcr-1* gene was carried in the chromosome of the clone ‘ST354/39 O1:H34’ in farm A and ‘ST117/48 O24:H4’ in farm C. The *mcr-1* gene was found at different positions of the genomes, each time next to the *pap2* gene (Fig. S6A, Fig. S6A). The *mcr-1* gene was detected in the same region as *papC* and *papG* allele II genes in the ‘ST117/48 O24:H4’ clone in farm C (Fig. S6A).

**Diffusion of ESBL-producing clones in calves and of *bla*_CTX-M_-carrying plasmids in clones**

The plasmidic localization of the *bla*_CTX-M_ and *mcr-1* genes was investigated using Southern blot hybridization, conjugation and genome sequencing. Thirty-one transconjugants carrying plasmid-borne ESBL-encoding genes and four transconjugants carrying plasmid-borne *mcr-1* gene were obtained, allowing the characterization of their genomic environment (Fig. S6A and Fig. S6B).

**Diffusion of ESBL-producing clones in calves and of *bla*_CTX-M_-carrying plasmids in clones**

There was a high diversity of ESBL-producing *E. coli* clones at the beginning of the fattening with 65.6% (21/32) of the clones detected during the first month (Fig. 3, Fig. S4A).

The presence of the *bla*_CTX-M-1_ gene in farm A was mostly due to the clone ‘ST10/466 O8:H32’ (28 isolates) carrying the plasmid IncF/F59:A-:B- and the clone ‘ST1850/918 O9:H10’ (22 isolates) carrying the plasmid IncI1/ST312 (Fig. 4B). They were widespread among the batch of calves, as 86.7% (13/15) and 80.0% (12/15) of calves excreted ‘ST10/466 O8:H32’ and ‘ST1850/918 O9:H10’, respectively (Fig. 3A). The presence of the *bla*_CTX-M-14_ gene in farm A was mainly due to four plasmids disseminated in five clones (encompassing 20 isolates) that were intermediate or inefficient spreaders among calves (Fig. 5A and Fig. 5B).

In farm B, only 18.2% of the clones (2/11) were isolated in different calves and were limited to two calves (Fig. 3B).

In farm C, the diversity of ESBL-producing clones isolated was the highest during the first month, with 8 clones detected at day 7 and 4 clones detected at day 21 (Fig. 3C, Fig. S4B). The presence of the *bla*_CTX-M-1_ gene was mostly due to the clone ‘ST329/919 O109:H16’ (26 isolates) carrying the plasmid IncI1/ST3 lineage 2 (Fig. 4B). This clone represented 72.2% (26/36) of the isolates carrying this gene and 35.1% (26/74) of all the isolates in this farm. It was found in 100% (14/14) of calves (Fig. 3C). The presence of the *bla*_CTX-M-14_ gene was mostly due to the clone ‘ST744/2 O101:H9’ (23 isolates) carrying an IncK plasmid. This clone represented 63.8% (23/36) of the isolates carrying this gene and 31.1% (23/74) of all the isolates in this farm. It was detected in 92.8% (13/14) of the calves. These two efficient colonizers carried plasmids that were locally inefficient spreaders, as no other clones in this farm were found to carry these plasmids. The *bla*_CTX-M-14_ gene-carrying IncF/F2:A-:B- plasmid had spread in five inefficient colonizers and a colonizer with intermediate efficiency, which represented 33.4% (12/36) of the isolates carrying this gene, and 16.2% of all the isolates in this farm (Fig. 5B). This plasmid was detected in 57.1% (8/14) of the calves.

**Diffusion of ESBL-producing clones and of *bla*_CTX-M_-carrying plasmids between farms**

The clones ‘ST10/2 O101:H9 clone2’, ‘ST167/2 O101:H9’, ‘ST301/917 O80:H2’ and ‘ST58/24 O9:H25’ were isolated in both farms B and C (Fig. 3). The clone ‘ST57/533 O119:H10’ was isolated in both farms A and C.

The mean number of SNPs between isolates of a clone detected in one farm was 9 (± 19), whereas the mean number of SNPs between isolates of a clone detected in several farms was 28 (± 16) (Fig. S1D).

**Diffusion and persistence of ESBL-producing clones over the fattening period**

The majority of the clones detected were isolated at a single time point in one calf, or simultaneously in two calves and were never isolated anymore (56.3%, 18/32 clones, Fig. 6). Eleven clones were detected in a larger number of calves and over longer periods.

**Diffusion of *mcr-1* gene in farms**

The IncX4 plasmid was found in the clones ‘ST10/466 O8:H32’, ‘ST10/2 O101:H9 clone3’ and ‘ST34/638 O68:neg’ in farm A (encompassing 27, 6 and 5 isolates resistant to colistin, Fig. 7). The *mcr-1* gene was never found on the same plasmid as the *bla*_CTX-M_ gene.

In four *mcr-1* gene carrying-clones, the gene was not detected in a subset of the isolates. This subset was either a minority of isolates (1/28 isolates for ‘ST10/466 O8:H32*’*, 2/7 for *‘*ST34/638 O68:neg’), or the majority of them (20/22 for ‘ST1850/918 O9:H10’, 2/3 for ‘ST10/383 O129:H11’).
