## Supplementary Tables S2 S3 for "Interplay between bacterial clone and plasmid in the spread of antibiotic resistance genes in the gut: lessons from a temporal study in veal calves"

Table S1. Summary of the origin and molecular characteristics of the ESBL-producing E. coli isolates and plasmids. Panel A represents the origin and molecular characteristics of the ESBL-producing E. coli isolates. Pulsed-field gel electrophoresis (PFGE) was performed on all putative ESBL-producing E. coli isolates using the restriction enzyme XbaI. Comparison of PFGE profiles was done among isolates to discriminate between ESBL-producing E. coli strains circulating in the farm and to select one representative of each PFGE profile for whole genome sequencing. Two isolates were considered to be related to different genotypes if their PFGE profiles differed from at least one band. Isolates having a different status regarding the presence of the mcr-1 gene compared to their group were additionally selected for genome sequencing. The clone ID and the sample code (the animal ID followed by the sampling rank) are also presented in the table. Panel B represents the donor strain and molecular characteristics of the plasmids carrying blaCTX-M genes and mcr-1 gene. The genome of transconjugants (TC) carrying plasmid-borne ESBL-encoding genes or mcr-1 gene were sequenced to characterize the molecular environment of these antibiotic resistance genes. Plasmids were identified using the PlasmidFinder database and reference genomes were found by blasting contigs against MaGe or Refseq databases.

Table S2. Posthoc Dunn’s test results, comparison of the number of copies of *bla*_CTX-M_ genes /g of feces (log_10_(copies/g)) estimated by quantitative PCR between the samples grouped according to the number of colonies on ESBL-producing *Enterobacteriaceae* selective medium. Quantification of *bla*_CTX-M_ group 1 and of *bla*_CTX-M_ group 9 gene copies was performed by qPCR on samples collected 7 days and 21 days after arrival in farms, and one by month for each calf.

| Comparison | Unadjusted p-value | Bonferroni adjusted p-value |
| --- | --- | --- |
| [No colony] – [0-100 colonies] | 1*10^-4^ | 1*10^-4^ ***** |
| [0-100 colonies] – [>100 colonies] | <10^-15^ | <10^-15^ *** |
| [No colony] – [>100 colonies] | <10^-15^ | <10^-15^ *** |

Table S3A. Phenotypic prevalence of antibiotic co-resistances in the ESBL-producing *E. coli* isolates (n= 173). Antibiotics are sorted in descending order according to their frequency and classes.

| Antibiotic classes | Antibiotics |  | Farm A  (n = 84) | |  | Farm B  (n = 15) | |  | Farm C  (n = 74) | |  | Total  (n=173) | |
| --- | --- | --- | --- | --- | --- | --- | --- | --- | --- | --- | --- | --- | --- |
|  |  |  | n | % |  | n | % |  | n | % |  | n | % |
| Cyclines | Tetracycline |  | 83 | 98.8 |  | 13 | 86.7 |  | 73 | 96.1 |  | 169 | 97.7 |
| Sulfonamides | Sulfafurazole |  | 82 | 97.6 |  | 15 | 100.0 |  | 67 | 90.5 |  | 164 | 94.8 |
|  | Streptomycin |  | 48 | 57.1 |  | 13 | 86.7 |  | 67 | 90.5 |  | 128 | 74.0 |
| Aminoglycosides | Kanamycin |  | 28 | 33.3 |  | 7 | 46.7 |  | 71 | 95.9 |  | 106 | 61.3 |
|  | Gentamicin |  | 21 | 25.0 |  | 0 | 0 |  | 4 | 5.4 |  | 25 | 14.5 |
| Diaminopyrimidines | Trimethoprim |  | 71 | 84.5 |  | 14 | 93.3 |  | 39 | 52.7 |  | 124 | 71.7 |
| Quinolones | Nalidixic acid |  | 20 | 23.8 |  | 6 | 40.0 |  | 36 | 48.6 |  | 62 | 35.8 |
|  | Enrofloxacin |  | 15 | 17.9 |  | 1 | 6.7 |  | 31 | 41.9 |  | 47 | 27.2 |
| Polymyxin | Colistin |  | 52 | 61.9 |  | 0 | 0 |  | 1 | 0.01 |  | 53 | 30.6 |
| Phenicols | Florfenicol |  | 23 | 27.4 |  | 4 | 26.7 |  | 4 | 5.4 |  | 31 | 17.9 |

Table S3B. Quinolone resistance mutations detected in the ESBL-producing *E. coli* isolates. Quinolone resistance point mutations were searched using the website “Center for Genomic Epidemiology” with standard parameters.

| Isolate ID | Farm | Clone name | Quinolone resistance level | GyrA S83 | GyrA D87 | ParC S80 | ParC A56 | ParC E84 | ParE I355 |
| --- | --- | --- | --- | --- | --- | --- | --- | --- | --- |
| 41086 | A | A ST10/2 O101:H9 *fimH54* clone3 | enrofloxacin | 1 | 1 | 1 | 0 | 0 | 0 |
| 41052 | C | A ST10/2 O101:H9 *fimH54* clone1 | enrofloxacin | 1 | 1 | 1 | 0 | 0 | 0 |
| 40772 | B | A ST10/2 O101:H9 *fimH54* clone2 | nalidixic acid | 1 | 0 | 0 | 0 | 0 | 0 |
| 41054 | C | A ST10/2 O101:H9 *fimH54* clone2 | nalidixic acid | 0 | 1 | 0 | 0 | 0 | 0 |
| 41299 | C | A ST744/2 O101:H9 *fimH54* | enrofloxacin | 1 | 1 | 1 | 1 | 0 | 0 |
| 40762 | B | B1 ST162/25 O8:H19 *fimH32* | enrofloxacin | 1 | 0 | 1 | 0 | 0 | 0 |
| 41048 | C | B1 ST949/21 O8:H16 *fimH121* | enrofloxacin | 1 | 1 | 0 | 0 | 1 | 0 |
| 41177 | C | C ST88/66 O21:H21 *fimH-* | enrofloxacin | 1 | 1 | 1 | 0 | 0 | 0 |
| 40808 | A | F ST354/39 O1:H34 *fimH58* | enrofloxacin | 1 | 1 | 1 | 0 | 1 | 0 |
| 40939 | A | F ST354/39 O1:H34 *fimH58* | enrofloxacin | 1 | 1 | 1 | 0 | 1 | 1 |
| 41058 | C | A ST167/2 O101:H9 *fimH215* | nalidixic acid | 0 | 1 | 0 | 0 | 0 | 0 |
| 40752 | B | A ST301/917 O80:H2 *fimH54* | nalidixic acid | 1 | 0 | 0 | 0 | 0 | 0 |
| 40769 | B | A ST301/917 O80:H2 *fimH54* | nalidixic acid | 1 | 0 | 0 | 0 | 0 | 0 |
| 41039 | C | A ST301/917 O80:H2 *fimH54* | nalidixic acid | 1 | 0 | 0 | 0 | 0 | 0 |
| 40892 | B | B1 ST58/24 O8:H25 *fimH34* | nalidixic acid | 1 | 0 | 0 | 0 | 0 | 0 |
| 41115 | C | E ST57/533 O119:H10 *fimH54* | nalidixic acid | 1 | 0 | 0 | 0 | 0 | 0 |
| 41533 | A | E ST57/533 O119:H10 *fimH54* | nalidixic acid | 1 | 0 | 0 | 0 | 0 | 0 |
| 41117 | C | G ST117/48 O153:H4 *fimH97* | nalidixic acid | 1 | 0 | 0 | 0 | 0 | 0 |
| 40759 | B | G ST117/48 O33:H4 *fimH41* | nalidixic acid | 1 | 0 | 0 | 0 | 0 | 0 |
